# Precise Generation of Conformational Ensembles for Intrinsically Disordered Proteins *via* Fine-tuned Diffusion Models

**DOI:** 10.1101/2024.05.05.592611

**Authors:** Junjie Zhu, Zhengxin Li, Zhuoqi Zheng, Bo Zhang, Bozitao Zhong, Jie Bai, Xiaokun Hong, Taifeng Wang, Ting Wei, Jianyi Yang, Hai-Feng Chen

## Abstract

Intrinsically disordered proteins (IDPs) play pivotal roles in various biological functions and are closely linked to many human diseases including cancer, diabetes and Alzheimer disease. Structural investigations of IDPs typically involve a combination of molecular dynamics (MD) simulations and experimental data to correct for intrinsic biases in simulation methods. However, these simulations are hindered by their high computational cost and a scarcity of experimental data, severely limiting their applicability. Despite the recent advancements in structure prediction for structured proteins, understanding the conformational properties of IDPs remains challenging partly due to the poor conservation of disordered protein sequences and limited experimental characterization. Here, we introduce IDPFold, a method capable of generating conformational ensembles for IDPs directly from their sequences using fine-tuned diffusion models. IDPFold bypasses the need for Multiple Sequence Alignments (MSA) or experimental data, achieving accurate predictions of ensemble properties across numerous IDPs. By sampling conformations at the backbone level, IDPFold provides more detailed structural features and more precise property estimation compared to other state-of-the-art methods. IDPFold is ready to be used in the elucidate the sequence-disorder-function paradigm of IDPs.

## Introduction

Intrinsically disordered proteins (IDPs) constitute a category of proteins with unstable structures under physiological conditions^1^. These proteins, accounting for over 40% of eukaryotic proteomes^2,3^, are involved in various biological functions including signal transduction, molecular recognition, and cell cycle regulation^4–6^. IDPs are also closely associated with various significant diseases, such as cancer, Parkinson’s disease, and acquired immunodeficiency syndrome (AIDS)^7–9^. Unlike structured proteins that possess one or a few stable conformations, IDPs exhibit transitions between multiple conformations with very low energy barriers, constantly fluctuating within a broad ensemble of structures under physiological conditions^10–12^. Consequently, deciphering the conformational ensemble of IDPs poses a significant challenge for experimental methods such as X-ray, cryo-electron microscopy and NMR^13–16^.

Currently, molecular dynamics (MD) simulation is the most commonly used and effective tool for sampling conformational ensembles^17,18^. By iteratively sampling target molecular system based on first principles, an estimation of conformational ensemble is obtained from simulation. However, MD simulation incurs high sampling costs, and ensuring thorough sampling of the ensemble is challenging. Additionally, commonly used force fields for IDPs, such as ff03CMAP^19^, a99SB-disp^18^, ESFF1^20^, still exhibit considerable errors in estimating local and global properties of IDPs during simulations^21,22^. Although we can adjust MD simulations to better align with the characteristics of IDPs using experimental data, only a few of these proteins have been experimentally characterized^23^. These limitations constrain the application of MD in predicting the conformational ensemble of IDPs.

On the other hand, numerous deep learning methods have been developed and widely applied in the field of structure prediction for structured protein, such as AlphaFold2 and ESMFold^24,25^. Simultaneously, in protein design tasks, various generative models, such as RFdiffusion and Chroma, have been utilized for generating diverse protein backbones^26,27^. Therefore, it is natural to consider whether deep learning methods can be employed for rapid and accurate prediction of conformational ensembles of IDPs. However, the structures associated with IDPs are currently very sparse. The PDB database contains > 220,000 structural entries^28^, while the Protein Ensemble Database only includes 553 protein ensemble data^29^. Moreover, IDPs often lack high-quality multiple sequence alignment (MSA) data. MSA has been proven effective only in functional studies concerning folded or bound states of IDPs, and provides little assistance in predicting more disordered conformations^30^. These challenges make it highly daunting to use deep learning for robustly predicting the conformational ensembles of IDPs. Although Janson et al. have previously worked on IDP conformation generation and proposed idpGAN for predicting IDP conformational ensembles, they mainly forced on generating coarse-grained IDP conformations and often suffers from over-sampling, leaving a gap to coarse-grained or even all-atom MD simulations^31,32^.

We introduce IDPFold here for predicting IDP dynamics directly from sequences based on a generative deep learning model. IDPFold utilized a protein language model to extract sequence information and further fed it into a structure generation module, enabling MSA-free conformation generation. To address the issue of insufficient data, we employed a hybrid dataset comprising crystal structures, NMR structures and MD trajectories for training IDPFold. The experimental structures enable the model to learn basic protein characteristics, while MD trajectories provide sufficient IDP structural data, ensuring accurate sampling on IDP systems. IDPFold generates IDP conformational ensembles at backbone level that is in better agreement with experimental observations than other state-of-the-art methods. IDPFold is able to sample conformational transitions from structured to disordered states, providing insights for studying the correlation between structures and functions of IDPs.

## Results

IDPFold employs a conditional diffusion model framework for generating protein conformational ensembles from sequences (Figure 1a). This framework involves a forward diffusion process where noise is gradually added to real protein structures, and a reverse diffusion process where a deep learning network is used for denoising. By integrating specific protein sequence features into the model during the reverse diffusion process, we can generate conformational ensembles for specific proteins using this architecture. The denoising network takes as input the sequence features extracted by ESM2 and consists of an initialization block and four denoising blocks (Figure 1b). The initialization module integrates the sequence features, noise scale, and noised structure, while denoising modules combine Invarient Point Attention (IPA) with traditional Transformers to capture the chain-like structure within proteins and the rotational/translational state of each residue (Figure 1c). For more details, please refer to the Materials and Methods section.

**Figure 1:**
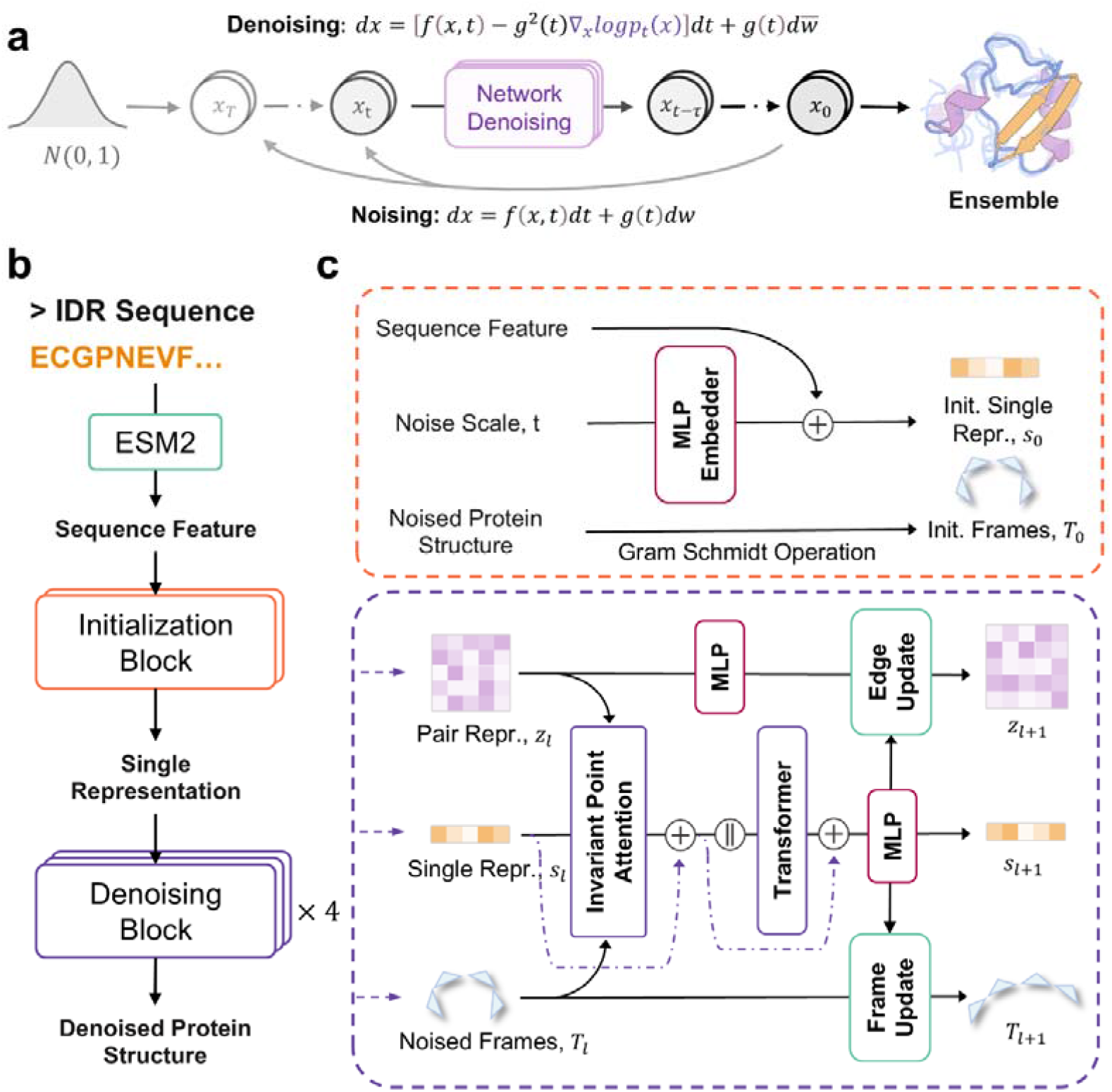
Architecture of IDPFold. **a**, Diffusion process for generating protein ensembles. **b**, Structure of denoising network. **c**, Detailed architecture of Initialization Block and Denoising Blocks.

### IDPFold reproduces global features of IDPs

We first evaluated the IDPFold-predicted ensembles at coarse-grained level, primarily focusing on the global characteristic of the predicted ensembles, specifically the radius of gyration (Rg). Rg is a critical physical quantity that reflects the global characteristic of IDPs with its magnitude positively correlating with overall protein looseness, making it an empirical measure to describe and distinguish IDPs from structured proteins^33,34^. We first referred to the evaluation by Tesei et al. on the coarse-grained force field CALVADOS 2 by calculating the Rg error of the IDPFold-predicted ensembles on their test set. After removing sequences presented in the training set, this test set contained 58 IDP systems. Among these proteins, the average Rg error of the IDPFold-predicted ensembles was − 3%, comparable to the performance of coarse-grained simulations reported by Tesei et al. (Figure 2a). We further examined the Rg distribution of predicted ensembles for each system and compared it with coarse-grained trajectories. The results demonstrated that the range of Rg distribution of IDPFold-predicted ensembles closely matched that of the coarse-grained simulations for most systems, indicating that IDPFold is capable of sampling a broad magnitude of IDP conformations similar to those observed in coarse-grained simulation trajectories and experimental values as well (Figure 2b, Figure S1).

**Figure 2:**
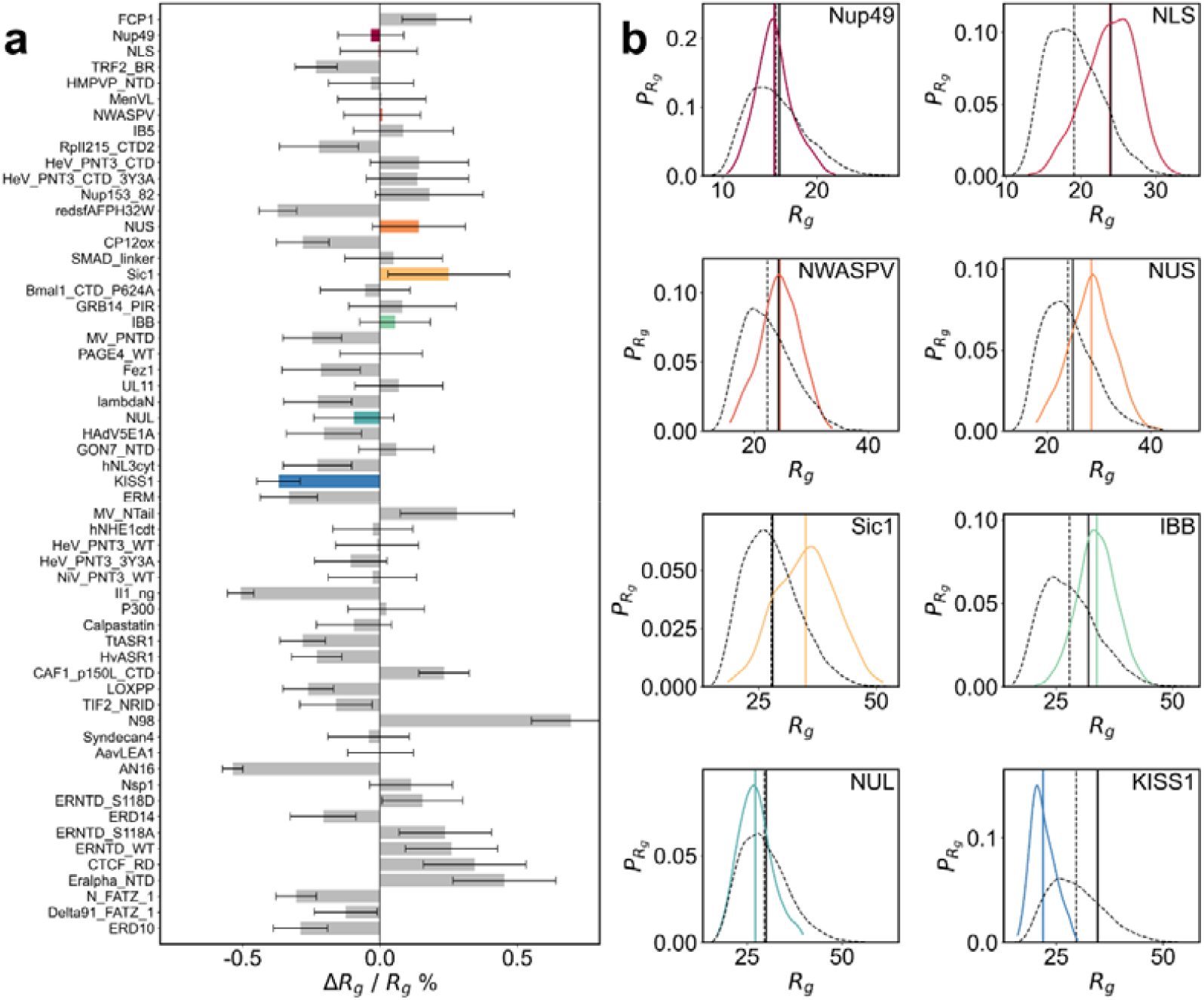
Comparison between the generated ensembles of IDPFold and coarse-grained MD simulation with CALVADOS 2. **a**, Relative error between IDPFold predicted and experimental radii of gyration on test set of CALVADOS 2. **b**, Rg distributions of IDPFold generated ensembles (colored) and CALVADOS simulations (black dashed). Experimental values are plotted as solid black lines.

### IDPFold captures the global distribution of IDP ensembles

To further assess the robustness of IDPFold, we collected 27 IDP systems which possessed rich experimental observation data and were not present in the training set (Table S1, S2). We compared the IDPFold-predicted ensembles with the simulation results from CALVADOS 2 and the predictions from the coarse-grained deep learning method idpGAN on these systems^23,31^. The result showcased that average Rg error of the IDPFold-predicted ensembles on this test set (*ε*_*Rg*_ = −0.06) was significantly smaller than that of idpGAN (*ε*_*Rg*_ = −0.12, with a paired t-test p-value of 0.02). The Rg error of the simulation trajectories was 0.02, which is better in absolute terms than both deep learning methods while without significant difference with IDPFold (a paired t-test p-value of 0.12). Compared to coarse-grained simulations, IDPFold tends to slightly underestimate Rg for IDPs, particularly for longer proteins. We hypothesize that this is partly due to the high conformational space complexity of long IDPs, making it more challenging to model their conformational ensembles. This is partially verified by the performance of the fine-tuned version of IDPFold, where the fine-tuning on MD trajectories corrected the disorder tendency in the generated conformations to a certain extent, while still underestimates the extent of extension for large systems. Overall, IDPFold tends to have a lower proportion of highly extended conformations in the predicted ensembles for large systems. This opens a future direction towards the improvements on the accuracy of global characteristic estimates by reweighting to increase the proportion of these extended conformations in the ensembles.

We next observed the ensemble distributions and main conformations generated by the three methods on specific cases. Here, we calculated the Rg-RMSD distribution of each ensemble using the initial conformations from coarse-grained simulations as a reference. The results showed that idpGAN exhibited over-sampling across all test systems and had significant deviations in estimating the main conformations. As a comparison, the ensemble distribution estimated by IDPFold was closer to coarse-grained trajectories, with its estimation of the free energy well positions more accurate than that of idpGAN (Figure 3a, c, e). Although the sampling range of IDPFold was smaller than that of coarse-grained trajectories, its estimation of the ensemble’s Boltzmann distribution was more precise than that of idpGAN.

**Figure 3:**
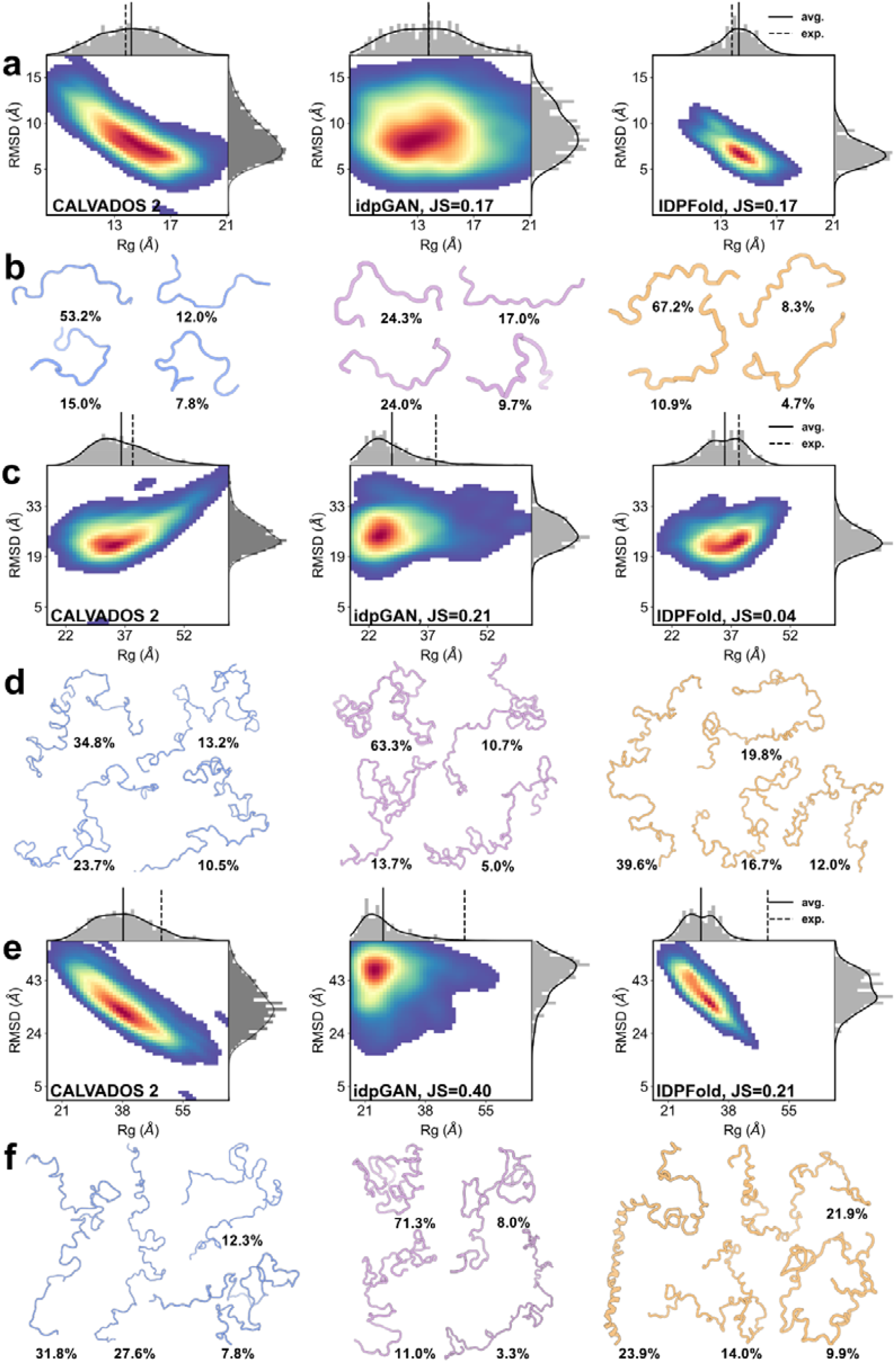
Comparing the generated ensembles of IDPFold and coarse-grained deep learning method idpGAN. **a, b**, Rg-RMSD distribution **(a)** and cluster centers **(b)** of predicted ensembles on Histatin5. **c, d**, Rg-RMSD distribution **(c)** and cluster centers **(d)** of predicted ensembles on Human Calpastatin. **e, f**, Rg-RMSD distribution **(e)** and cluster centers **(f)** of predicted ensembles on β Synuclein.

From the clustering results, the main conformations and their proportions sampled by IDPFold across the three systems were closer to the simulation results than idpGAN. In the Histatin5 protein, both IDPFold-predicted ensembles and the simulation trajectories had an extended conformation as the main structure, whereas the main conformation predicted by idpGAN was in a semi-folded state (Figure 3b). In the two larger systems, Human Calpastatin and β Synuclein, the issues with idpGAN were even more pronounced, as it predicted collapsed main conformations, which led to a severe underestimation of the average Rg of these two proteins (Figure 3d, f). This indicates that idpGAN’s estimation of ensemble distribution is inaccurate. On the other hand, IDPFold-predicted ensembles more accurately captured the shapes of the main conformations. Although IDPFold underestimated the Rg on β Synuclein due to overestimating the helical tendency in the structure, its estimates of the proportions of extended and folded conformations were closer to the simulation results.

### IDPFold outperforms conventional all-atom simulation

A major advantage of IDPFold compared to the aforementioned coarse-grained simulations is its capability to generate protein backbones, allowing us to better understand the dynamic properties of proteins at higher precision. Thus, we conducted all-atom molecular dynamics (MD) simulations on 27 IDP systems aforementioned in the test set to assess the quality of the local features in the IDPFold-predicted ensembles. We used the ESFF1 force field which was specifically parameterized for IDPs, and solvent model OPC3-B^20,22^. To demonstrate the convergence of our simulation, we performed three sets of repeated simulations on 10 of these proteins, recording the distribution and mean of experimentally observed physical quantities across the parallel trajectories. The results of the convergence analysis are shown in Figures S2. Subsequently, we compared the IDPFold-predicted ensembles with the results from the all-atom MD simulations.

We first examined the distributions of bond lengths, bond angles, and dihedral angles predicted by IDPFold. In terms of bond length and bond angle distributions, the ensemble predicted by IDPFold closely resembles those observed in the MD trajectories, indicating that the model has effectively learned the arrangement of side chains in protein residues, accurately predicting the distances and arrangements between adjacent atoms (Figure 4a). Additionally, the distribution of *Ω* angles in the conformations generated by the model resembles that observed in MD trajectories showcased a peak around 180°, suggesting that the local peptide bond plane conformations generated by the model are stable and align with general protein characteristics (Figure 4b). Moreover, the Ramachandran plot of *φ* − *ψ* angles shows that the backbone dihedral angles of generated conformations predominantly fall within reasonable regions. The density estimation across various regions also closely approximate those observed in MD trajectories (Figure 4c-e). Specifically, with IDPFold fine-tuned on IDRome data, there is a notable improvement on the probability of the ppII region in the upper left corner compared to the untuned version, showing a distribution closer to MD trajectories. This indicates that the fine-tuning enables the model to generate more disordered structures, capturing the intrinsic biases of target proteins more precisely.

**Figure 4:**
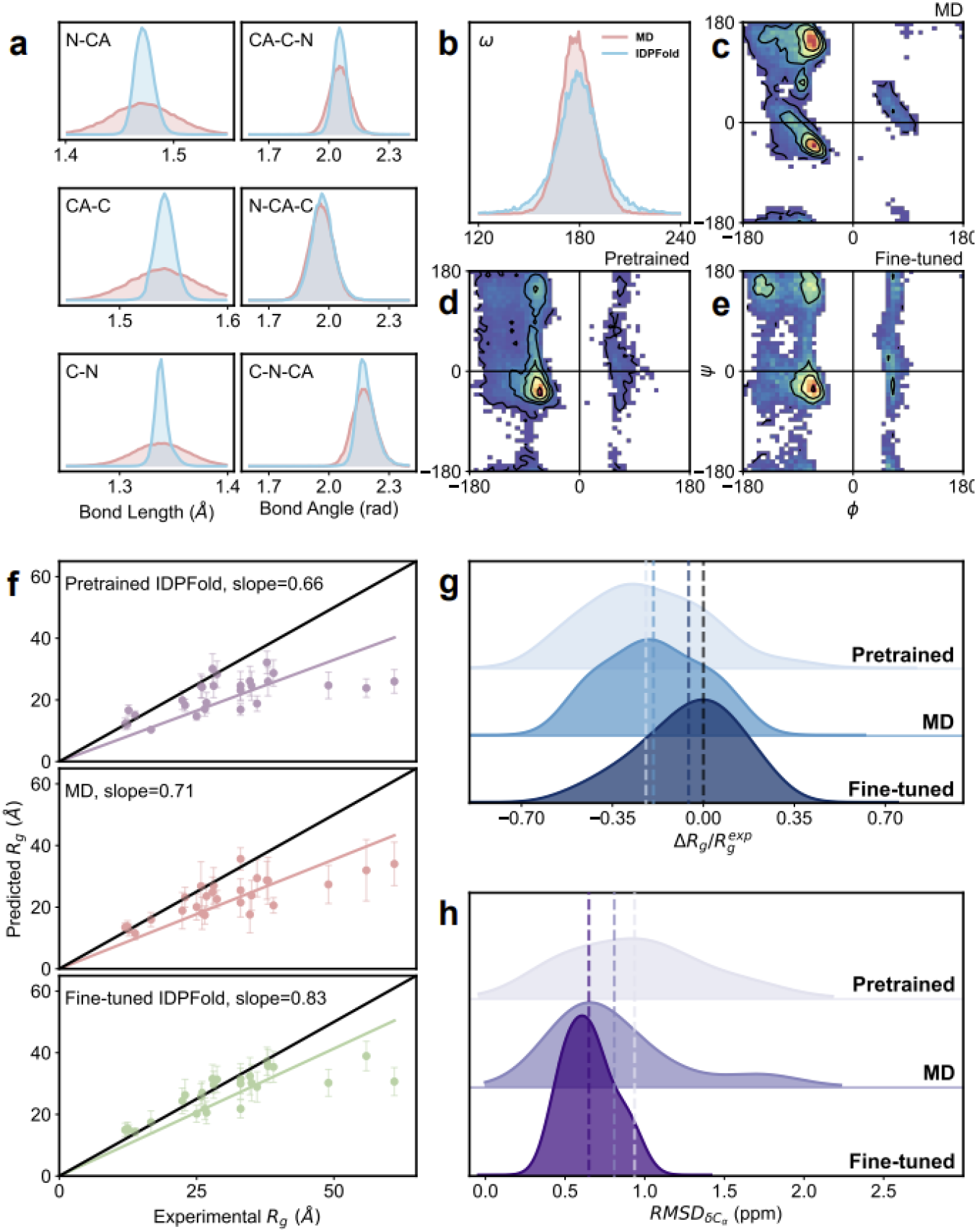
Comparison between the generated ensembles of IDPFold and all-atom MD simulation with ESFF1. **a-c**, Bond length **(a)**, bond angle **(b)** and omega angle **(c)** distributions of IDPFold generated ensembles (colored) and MD simulation (black). Lines in orange denotes inter-residue bond properties. **d-f**, Ramachandran plots of MD simulations **(d)**, pretrained IDPFold generated ensembles **(e)** and finetuned IDPFold generated ensembles **(f). g, h**, Ensemble average Rg error **(g)**, RMSD on ensemble average *C*_*α*_ chemical shift **(h)** of MD trajectories (middle), pretrained (upper) and fine-tuned (lower) IDPFold against experimental values. **i**, Regression plot between experimental Rg values and predicted Rg values for MD simulation and IDPFold. The intercept was set to zero.

We next assessed ensemble average Rg of the generated ensembles (Figure 4f, g). We calculated the average error and standard deviation relative to experimental Rg values for conformational ensembles generated by IDPFold. Those generated by IDPFold pretrained on experimental structures significantly underestimated Rg for most IDPs (average error of −0.22), indicating a clear bias towards structured protein in prediction. This observation aligns with the high proportion of structured regions in experimental structures, lacking representation of natural disordered regions. However, the model significantly improved its Rg estimation after fine-tuning on MD trajectories, with an average error of only −0.06. Paired t-test demonstrates a significant improvement over all-atom simulation (average error of −0.19) with *p* = 3.57 ×10^−4^.

IDPFold not only accurately captured the ensemble average Rg of IDPs but also exhibited considerable diversity in generated conformations comparable to MD simulations (Figure S3, S4). Particularly, for IDP systems with a higher structured tendency, such as ACTR and drkN SH3, the pretrained IDPFold exhibited minimal variance and mean Rg in generated structures, indicating limited diversity and excessive compactness. As a comparison, the fine-tuned model showcased average Rg values closer to experimental observations with deviations comparable to MD trajectories (*p* = 0.22 in paired t-test), suggesting both a higher diversity in generated conformations and better alignment with experimental states.

Additionally, we evaluated the local features of generated ensembles using chemical shifts as the target physical quantity to measure how generated ensembles differ from experimental observations (Figure 4A, Figure S5). Specifically, we computed *C*_*α*_ secondary chemical shifts of generated conformations and the Root Mean Square Deviation (RMSD) between ensemble average and experimental values. *C*_*α*_ secondary chemical shifts can reflect the secondary structure propensity of a protein system, where positive values indicate a higher helical propensity in certain region, negative values indicate a tendency to form strands, and values close to zero indicate disordered regions. The ensembles generated by fine-tuned IDPFold are close to experimental observations, showing an average RMSD of 0.53. Pairwise t-tests revealed significant improvement over MD simulations in fitting chemical shifts (*p* = 0.03), whereas pre-trained IDPFold performed worse than MD in fitting chemical shifts.

### IDPFold predicts IDP ensembles comparable to long MD simulations

Additionally, we collected trajectories used by Robustelli et al. in testing the a99SB-disp force field, along with other force field trajectories for comparison^18^. We analyzed the differences in performance between IDPFold and four all-atom molecular dynamics simulations on seven proteins that overlapped with our test set. These 30-microsecond trajectories were confirmed to have converged. We primarily focused on the Rg error and the RMSD of *C*_*α*_ and *C*_*β*_ secondary chemical shifts (Figure 5a).

**Figure 5:**
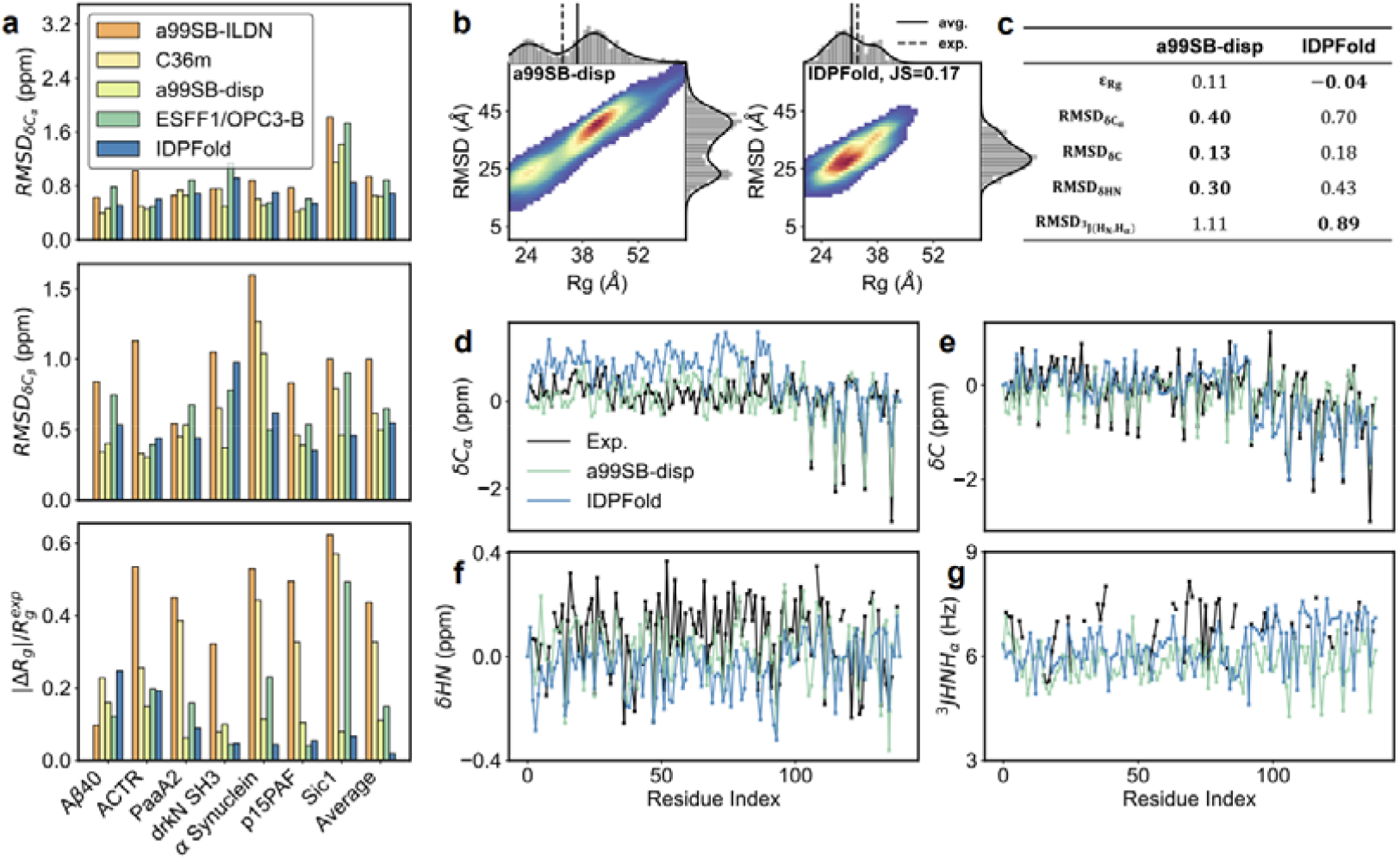
Comparison between the performance of IDPFold and that of all-atom MD simulations with different force fields. **a**, Accuracy of IDPFold in estimating local features (*C*_*α*_, *C*_*β*_ chemical shifts) and global feature Rg compared to MD simulations. **b**, Rg-RMSD distribution of IDPFold generated ensemble compared to MD simulation with a99SB-disp on α Synuclein. **c**, Errors between generated ensembles and experimental observations for a99SB-disp and IDPFold on α Synuclein. **d-g**, Local Features including *C*_*α*_ chemical shifts **(d)**, *C* chemical shifts **(e)**, *HN* chemical shifts **(f)** and ^3^*J*(*HN, H*_*α*_) **(g)** of IDPFold generated ensembles and MD simulation with a99SB-disp on α Synuclein.

In terms of chemical shift accuracy, the error in the IDPFold-predicted ensembles reached a level comparable to a99SB-disp, surpassing traditional force fields such as a99SB-ILDN and CHARMM36m^17,35^. Furthermore, IDPFold demonstrated significantly better performance in Rg error compared to all four force fields, indicating that IDPFold accurately captures the global characteristics of IDPs while achieving a comparable level of precision in local structure prediction as current state-of-the-art force fields. Using *α* Synuclein protein as an example, we illustrated the differences between the ensembles sampled by IDPFold and MD trajectories from the a99SB-disp force field. We calculated the Rg-RMSD distributions from both methods and found that while a99SB-disp sampled a wider range of conformations, capturing more extended states, IDPFold predominantly sampled more compact conformations (Figure 5b). In terms of Cα chemical shifts, we observed that IDPFold significantly overestimated the helical propensity in the N-terminal region of *α* Synuclein, while achieving a similar level of accuracy as a99SB-disp in the C-terminal region (Figure 5c, d). This also explains the tendency of IDPFold to sample more compact structures. For C and HN secondary chemical shifts, RMSDs of IDPFold were slightly larger than those of a99SB-disp, while in ^3^*J*(*HN, H*_*α*_) scalar coupling, IDPFold slightly outperformed a99SB-disp. Overall, we conclude that IDPFold’s characterization of IDP local features is on par with traditional force fields, and its sampling of α Synuclein takes around 20 minutes, significantly faster than the hundreds of hours required by traditional MD simulations. This demonstrates the power of IDPFold to serve as a complementary tool to traditional force field sampling which might pave the way for the exploration of macromolecular systems on a considerable time scale.

### IDPFold surpasses current state-of-the-art deep learning methods

Lastly, we compared IDPFold with existing deep learning methods on the test set, using experimental observations as the primary evaluation metric to establish a benchmark for ensemble prediction methods (Table 1, Figure S6). This benchmark included three coarse-grained methods and four methods with backbone-level or higher accuracy, in which AlphaFlow of the best performance (AF-PDB-base) is presented (Table S3). For local features, IDPFold achieved the best performance in most of the metrics among the backbone-precision methods (Figure S7-S13). In terms of global characteristic like Rg, IDPFold was slightly outperformed by the coarse-grained force field CALVADOS 2. We also evaluated the validity of the conformations generated by various deep learning methods (as defined in the materials and methods section). The results showed that all existing deep learning methods generated conformations with high validity. However, we observed a slight decrease in validity of IDPFold-generated ensembles after fine-tuning, likely due to the greater local structural fluctuations in the trajectory data used for fine-tuning compared to experimental structures which led to more pronounced local fluctuations in the generated conformations (Figure S14). Overall, IDPFold-predicted ensembles outperformed existing deep learning methods in all experimental observations and were comparable to or even better than traditional MD simulations.

**Table 1.** Benchmark on IDPFold and other methods. Bold values denote the best.

| Methods | Validity | $\epsilon_{Rg}$ | $RMSD_{\delta C_{\alpha}}$ | $RMSD_{\delta C_{\beta}}$ | $RMSD_{^3J(H_N, H_{\alpha})}$ | $RMSD_{RDC_{NH}}$ |
| --- | --- | --- | --- | --- | --- | --- |
| CALVADOS 2<sup>36</sup> | - | **0.02** | - | - | - | - |
| idpGAN<sup>31</sup> | 0.91 | -0.12 | - | - | - | - |
| idpSAM<sup>37</sup> | 0.95 | -0.55 | - | - | - | - |
| MD 3*1μs<br>ESFF1+OPC3-B | - | -0.19 | 0.81 | 0.64 | **0.96** | 3.37 |
| AF-cluster<sup>38</sup> | **0.99** | -0.12 | 0.74 | 1.30 | 1.81 | 3.84 |
| AlphaFlow<sup>39</sup> | 0.97 | -0.24 | 0.71 | 1.16 | 1.26 | 4.02 |
| IDPFold | 0.95 | -0.06 | **0.65** | **0.53** | 1.01 | **3.27** |

## Discussion

In this study, we developed IDPFold, a deep learning-based tool for predicting IDP dynamics conformation ensembles. IDPFold adopts a conditional diffusion model architecture to perform an end-to-end conformational generation, integrating protein language model for sequence feature extraction and DenoisingIPA module for conformation denoising. Through a two-stage training strategy on experimental data and MD trajectories of IDPs respectively, IDPFold efficiently and accurately samples IDP dynamics conformational ensembles. Furthermore, we established a benchmark of conformation sampling methods on 27 protein systems that contain IDRs. IDPFold precisely capture the overall compactness and local secondary structure features of IDPs, with predicted ensembles exhibiting features with values closer to experimental observations compared to both existing MD-based and deep-learning methods. Additionally, IDPFold can sample conformational changes from structured to disordered states in proteins, demonstrating a wide sampling range and high efficiency in ensemble sampling. This capability provides important insights for studying the conformational changes and functions of IDPs.

IDPFold achieves accuracy levels comparable to, or even higher than, traditional MD simulations in estimating IDP conformational ensembles with its sampling process not restricted by energy barriers. This is not only essential for studying IDP conformations but also holds tremendous promise for dynamic proteins like allosteric proteins and enzymes, which undergo large-scale conformational changes during functional processes. Although this work focused on training the model for IDPs and direct prediction performance on allosteric protein ensembles might not be optimal, the methodology would likely to be transferred on diverse categories of proteins, enabling more accurate estimations of conformational transitions in these functionally important proteins (Figure S15, Movie S1).

Although IDPFold demonstrates robustness and precision in predicting conformational ensembles, its inference time is relatively long compared to some of the existed deep learning methods with an average time of about 20 minutes to sample an entire system. While this speed is much faster than conventional MD simulations, there is room for further improvement on model efficiency. To achieve higher sampling accuracy and more precisely protein backbone features, we employed a more complex network architecture and longer diffusion steps which might well sacrifice sampling efficiency. To enhance the inference efficiency of the model, optimizations can be explored such as reducing the number of diffusion steps in the diffusion model and optimizing transformer components by replacing them with architectures that are more efficient in terms of time and space. Additionally, in this study, we only trained and tested IDPFold on single-chain proteins and did not evaluate it on protein complexes or multi-chain proteins. In future work, we aim to extend the model to these areas to efficiently sample and interpret how molecules dynamically interact with each other.

Conformational ensemble prediction represents a significant frontier in protein structure research following static structure prediction. Accurate characterization of protein conformational ensembles can help us understand the dynamics of proteins and crucial conformational changes they undergo when binding to substrates or undergoing biochemical reactions under physiological conditions. Although deep learning-based tools are not yet widely adopted and most dynamic analyses still relying on MD simulations, our work suggest that properties predicted using deep learning methods might offer higher accuracy than MD simulations, potentially serving as complementary or even alternative approaches to MD simulations. However, current deep learning-based conformational ensemble prediction methods still face several challenges, such as inaccurate estimation of conformational free energies, less robustness compared to traditional methods, and the inability to capture temporal autocorrelation between conformations (i.e., inability to capture dynamic features).

Several studies have indicated that the single-point energy estimation of molecular force fields can be used to train diffusion models, allowing models to learn more comprehensively about conformational space in cases where structural data are scarce^40,41^. This training approach, which utilizes force field energies rather than structural data, enables more accurate capture of conformational space distribution and greater robustness. However, this training strategy also makes the model sensitive to empirical parameters of the force field, demanding high precision in molecular force field accuracy. Additionally, there are works based on flow models or score-matching models aiming to fit the dynamic characteristics of MD trajectories, which might be a solution to the current inability of generative models to capture temporal autocorrelation between conformations^42^.

Apart from the challenges of sparse training data and limited model representation capabilities, there is another significant issue in conformational ensemble prediction tasks: the scarcity and lack of uniformity in evaluation metrics. While we collected a set of experimental observations as a gold standard for this study, a vast number of protein systems lack experimental annotations, making it challenging to assess the generative performance of computational models on these systems. Therefore, we believe that constructing confidence metrics for conformational ensemble generation akin to pLDDT for protein structure prediction is also promising for the evaluation and improvement of protein conformational ensemble models.

## Materials and Methods

### Datasets

The data for training IDPFold consist of three components: high-quality crystal structures collected from the PDB dataset, conformations ensembles resolved by NMR and molecular dynamics (MD) simulation dataset of IDPs obtained through extensive back-mapping and energy minimization on coarse-grained trajectories.

For crystal structures, we referenced trRosetta and gathered a total of 15,051 X-ray resolved structures with resolutions ≤ 2.5Å and sequence redundancy ≤ 30%. These structures provide fundamental protein characteristics for the model, such as bond lengths, bond angle distributions and chirality of *C*_*α*_. However, crystal structures typically depict stable conformations of proteins, i.e., structured conformations. Training the model solely on crystal structures would significantly underestimate the disorder tendency of IDPs and result in low diversity in conformation generation for individual protein sequences. Therefore, we additionally collected 12,339 NMR resolved protein conformational ensembles from the PDB. We filtered these NMR entries based on 30% sequence similarity both internally and against crystal structures, resulting in 539 systems comprising a total of 10,454 structures. By blending crystal structures with NMR ensembles, we obtained a combined total of 25,495 experimental structures for the initial phase of IDPFold training. These structures were well-defined and exhibited a higher structured tendency, ensuring that the model captures the local physical characteristics of proteins effectively.

For the IDP trajectory data, we obtained large-scale coarse-grained simulation data from IDRome that recorded most IDRs in human proteome^23^. These coarse-grained simulations accurately capture global features of IDPs (and IDRs), such as average radius of gyration (Rg) of conformational ensembles, but they only retain *C*_*α*_ atoms, whose resolutions fail to meet the requirement for model training. Therefore, we selected all systems with lengths exceeding 256 residues, totaling 3,880 systems. Choosing larger systems is primarily because these long disordered segments encompass most sequence and structural characteristics found in smaller systems, and training the model on larger systems can help enhance its generalization ability. For these systems, we first used pdbfixer to restore structures from coarse-grained to all-atom and then performed 100 steps of energy minimization with ff14SB^44^ force field for all protein structures. A total of 77,600 optimized all-atom conformations was used for the second phase of IDPFold training, these conformations globally exhibit more disordered and can aid the model in learning distinctive conformational features of natural IDPs.

Furthermore, data used to assess the performance of IDPFold generation included 27 IDP systems that are fully described by experiments and are not present in the training set. We conducted 1*μs* all-atom MD simulations for these systems with the IDP-specific force field ESFF1 and solvent model OPC3-B^20,45^. These all-atom simulation trajectories described the dynamics and thermodynamic characteristics of IDPs, which are suitable for evaluating the quality of IDPFold-generated conformational ensembles.

### Formulation of IDPFold

#### Diffusion Modeling on Protein Structure

To enable IDPFold to capture Boltzmann distribution of protein conformations at equilibrium state, we employ Score-Based Generative Modeling (SGM) to learn the probability distribution from protein structure data. SGM can be represented by a diffusion process *x*_*t*_ ∈ ℝ^*n*^ defined by stochastic differential equation (SDE)^46^. The forward diffusion process is characterized by the following equation:

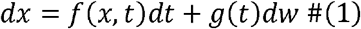

where *t* ∈ [0, *T*] is a continuous index and *w* ∈ ℝ^*n*^ is the standard Wiener process (a.k.a., Brownian motion). *f*(*x, t*) ∈ ℝ^*n*^, is a vector-valued function called the drift coefficient, and *g*(*t*) ∈ ℝ is a scalar function called the diffusion coefficient. Then, the corresponding backward diffusion process, or denoising process can also be defined by SDE^47^:

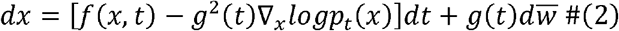

where 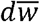 is a standard Wiener process as continuous time *t* flows backwards from *T* to 0, and *dt* is an infinitesimal negative time step. *x*_*t*=0_ here represents ground truth data, or the protein conformations, while *x*_*t*=T_ is sampled from Gaussian distribution. Therefore, by solving the backward SDE process, we can sample diverse protein conformations that obey Boltzmann distribution from a Gaussian distribution. In equation 2, each term except the score of *x*, ∇_*x*_ *logp*_*t*_ (*x*), is solvable. Therefore, we only needed a score-matching network to fit ∇*x logp*_*t*_ (*x*) at each time step to achieve the purpose of generating conformations.

Based on the above-mentioned standard formulation of SGM, we further required that the entire conformation generation process should be SE(3)-equivariant, i.e., diffusion process and network transformation were not sensitive to global rotation and translation of protein structures. SE(3)-equivariance can be described by the following equation:

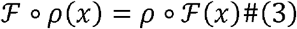

where ℱ denotes data transformations like network prediction and diffusion process, *ρ* represents global rotation and translation. Typically, a protein conformation *x* is characterized by Cartesian coordinates *c*_*i*_ ∈ ℝ^3^,1 ≤ *i* ≤ *N*, where *N* denotes number of atoms. However, data transformations on Cartesian coordinates is computationally intensive and does not easily satisfy SE(3)-equivariance. Therefore, we adopted backbone frame parametrization, representing the protein conformation *x* as *T*_*j*_ ≔[*R*_*j*_, *v*_*j*_], 1≤ *j* ≤ *n*, where *n* denotes the number of residues. Each backbone frame includes a 3 × 3rotation matrix *R*_*j*_ ∈ *SO*(3) and a translation vector *v*_*j*_ ∈ ℝ^3^. A frame *T*_*j*_ can represent Euclidean transformation for each atom in residue *j* from local coordinates *c*_*local*_ to global coordinates *c*_*glocal*_ as *c*_*glocal =*_ *T*_*i*_ ∘ *c*_*local*_ ≔ *R*_*i*_ *c*_*local*_ + *v*_*i*_. Following the approach described in FrameDiff^41^, we independently handle the rotation matrix *R*_*j*_ on the *SO*(3) manifold and the translation vector *v*_*j*_ in ℝ^3^during. the diffusion process, formulated as follows:

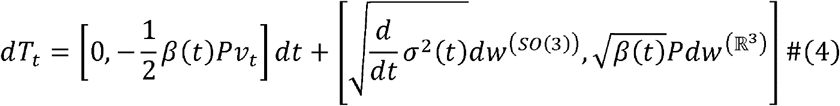

Where *β*(*t*) and σ(*t*) control the scale of noise during the diffusion process,*W*^ℳ^ denotes Brownian motion defined on manifold ℳ. and *P*:ℝ^3*n*^ → ℝ^3*n*^ is used for removing the center of mass. During the forward diffusion or noising process, the addition of noise on rotation matrices is determined by the noise kernel *p*_*t*|0_ (*R*_*t*_| *R*_0_), which is obtained from an isotropic Gaussian distribution on the *SO*(3)manifold.

This distribution is formulated as:

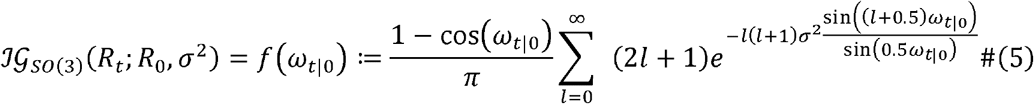

Here, 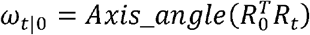 is the axis-angle transformed representation of the composed rotation matrix 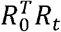. As for the translation vector, its noise addition process follows an Ornstein-Uhlenbeck process, also known as VP-SDE. The noise kernel for the translation vector is relatively straightforward, converging ultimately to 𝒩(0, *I*) as shown in the following equation.

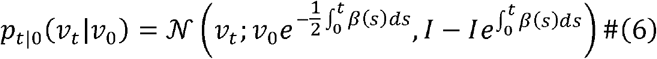

#### Network Design and Training

To achieve the goal of predicting conformational ensembles from sequence, we devised a sequence-conditioned score-matching model *s*_*θ*_(*x*_*t*_, *x, seq*)for denoising processes. As the diffusion process is designed to be strictly SE(3)-equivariant, network transformations should evidently preserve this property. Therefore, we adopted a variant of structure module from AlphaFold2 (Figure 1A) for updating backbone frames^24,32^. Here, we employed the Point Invariant Attention (IPA) mechanism to capture interactions and relationships between nearby residues, followed by a Transformer to learn global features and long-range interactions. This architecture has been demonstrated in previous research to promote training and generation of high-quality protein conformations. The aforementioned network design requires three inputs foreach layer: a one-dimensional vector representation *s*_*l*_, pairwise feature representation *z*_*l*_, and the set of rotation and translation updates *T*_*l*_. We utilized ESM2-650M to extract protein sequence features, concatenated with residue position encoding and time encoding represented by trigonometric functions, to form the initial one-dimensional vector representation *s*_*0*_. Pairwise feature representation *z*_*0*_. is derived from *s*_*0*_ based on relative positional encoding. After each layer of IPA-Transformer transformation, we updated the one-dimensional vector representation through a fully connected network and subsequently updated the pairwise features via cross product.

The objective of score-matching networks differs from conventional neural network training goals. It does not aim to fit protein conformations directly but rather the scores of data perturbed to a certain degree, i.e., fitting the distribution of perturbed data. To measure how well the predicted scores fit actual distribution, we computed the DSM loss as follows:

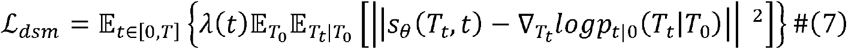

To ensure that the DSM loss at all time steps *t* results in a perfect fit score of 1, ensuring equal contribution of each time step to the loss function, we set the weights as follows:

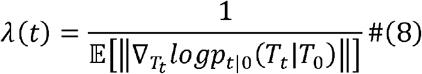

Additionally, to ensure the model learns detailed features of protein structures, apart from the DSM loss on rotation and translation matrices, we also incorporate mean square error (MSE) supervision for the positions of backbone atoms and differences on the distance matrix for samples with fewer forward diffusion steps 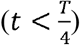. Therefore, the complete network training loss function can be represented as:

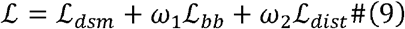

where *ω*_1 =_ *ω*_2_ = 0.25 control the weight of conformation quality loss.

During the training process, with a maximum time step *T* = 1.0, we optimize the DenoisingIPA using an Adam optimizer with the learning rate of 10^−4^^48^. For the translation vector part of the network training, we employ a linear noise strategy within the VP-SDE framework, while for the rotation matrix part, we use a logarithmic noise strategy within the VE-SDE framework, as shown in the following equation^46^:

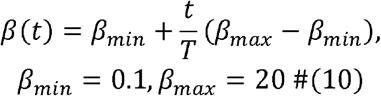

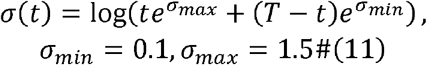

#### Implementation Details

The training of IDPFold consists of two stages: pre-training on experimental structures and fine-tuning on MD trajectories. In Table 2, we present the model hyperparameters utilized during the 2 training stages:

**Table 2.**
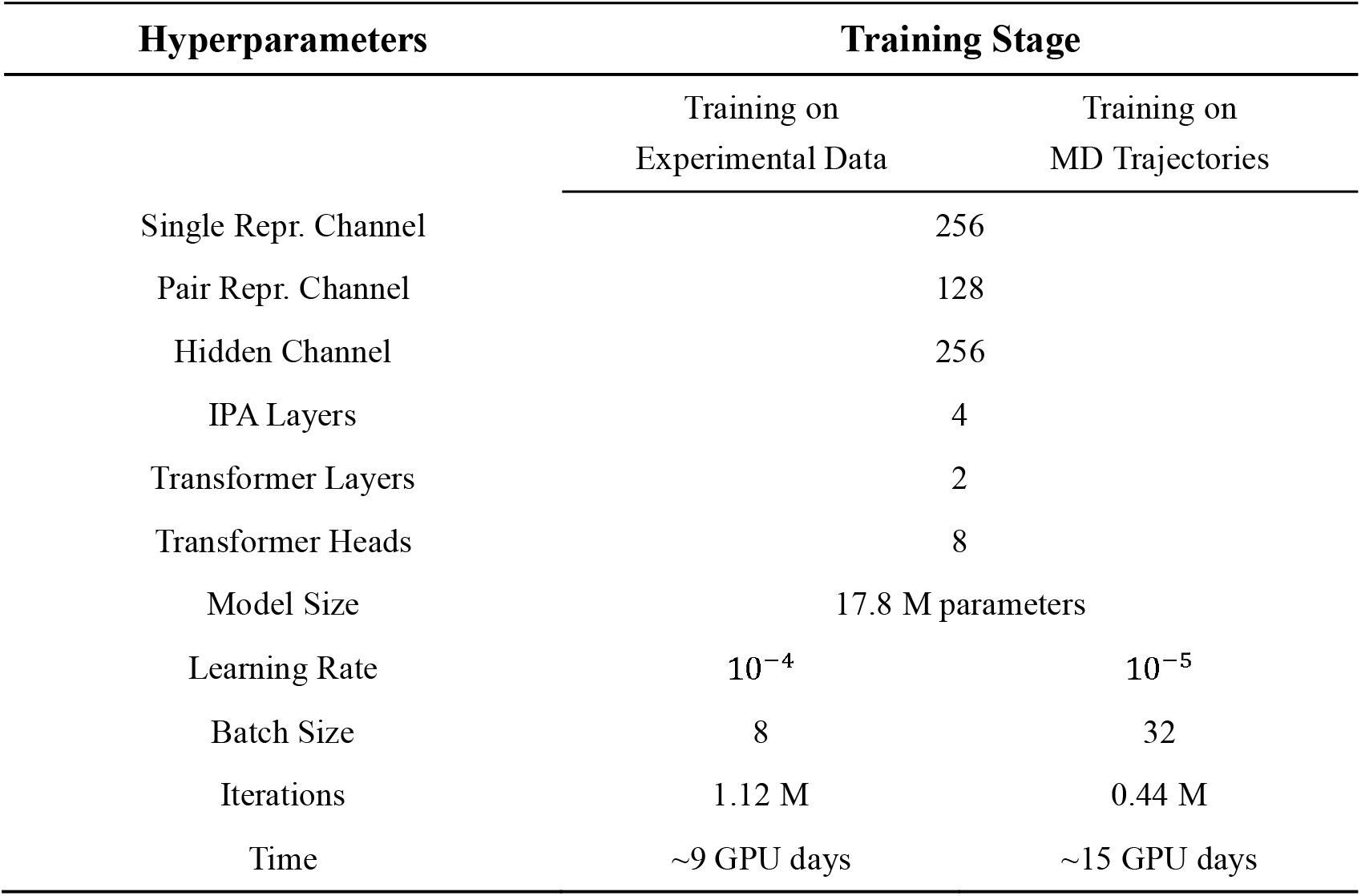
Hyperparameters and training details of IDPFold.

### Evaluation Metrics and Analysis Tools

We evaluated the quality of IDPFold-predicted IDP conformational ensembles from two perspectives: local features of the generated structures and global features of the conformational ensembles. For local features, biotite is used to compute inter-residue bond lengths and bond angles of all generated conformations^49^. We also analyzed the distribution of backbone dihedral angles to assess the model’s prediction accuracy regarding *C*_*α*_ chirality and secondary structure. Additionally, we used mdtraj to calculate scalar coupling between HN and *H*_*α*_, employed SPARTA+ to compute chemical shifts, and then performed ensemble averaging^50,51^. These analyses helped determine whether the local environment of the protein backbone aligns with experimental observations.

For global features, we calculated Rg of the generated conformations and compared the ensemble average with experiment results. Additionally, we calculated RMSD of generated conformations against the initial structures utilized in MD simulation in order to construct Rg-RMSD space. Through clustering the generated conformations and projecting them onto Rg-RMSD space, we aim to discern the diversity of generated conformations and assess the model’s learning efficacy in capturing the Boltzmann distribution information from MD trajectories in the training dataset. MMTSB toolset is applied for conformation clustering.

Due to the fact that most current work on generating protein conformational ensembles from sequences adopts a coarse-grained representation, where only *C*_*α*_ coordinates are generated, we first fixed the generated structures with pdbfixer and ran a 100-step energy minimization. During minimization we added restraints on *C*_*α*_ to make sure the optimized backbones did not differ from the original ones too much. This process cost about 6 seconds per conformation. Then we evaluated Validity and Fidelity and compared the performance of IDPFold with previous methods.

Validity assesses whether the generated conformations contain unreasonable *C*_*α*_ distances. In a protein structure, due to van der Waals interactions between atoms, *C*_*α*_ distances should not be too close. Additionally, because neighboring *C*_*α*_ in the protein backbone are connected by *C −N* with specific bond lengths, *C*_*α*_ distances should not be excessively far apart. Therefore, following the approach of Str2Str, we defined a reasonable range for *C*_*α*_ distances as follows:

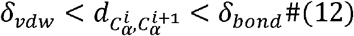

where *δ*_*vdw*_ = 2 ×1.7− 0.4 is defined as the sum of two *C*_*α*_ van der Waals radius minus an acceptable overlap distance of 0.4^52^.*δ*_*bond*_ is taken as the maximum *C*_*α*_ distance observed in MD trajectory of each test system. Since a reasonable range is defined, we could define validity as proportion of valid conformations. A higher validity indicates a lower probability of mis-estimated *C*_*α*_ distances, thus demonstrating better performance.

Fidelity refers to how well the model-generated ensembles match experimental observations. We selected Rg, *C*_*α*_ and *C*_*β*_ secondary chemical shifts, J-coupling constants between *HN* and *H*_*α*_, and backbone N-HN RDCs as target physical quantities to measure the fidelity of models, calculating the errors between model-generated conformational ensembles and experimental observations using the following formula:

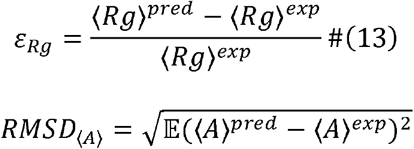

where ⟨*Rg*⟩ denotes the ensemble average Rg, ⟨*Rg*⟩^*pred*^ is the generated ensemble average Rg while ⟨*Rg*⟩^*exp*^ is experimental observation. Similarly, ⟨*A*⟩^*pred*^ and ⟨*A*⟩^*exp*^ denotes ensemble average physical quantity (e.g., chemical shifts and J-couplings) and experimental observation respectively. Using the above definitions, we calculated, *ε*_*Rg*_, 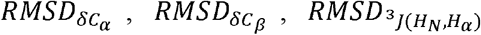, and 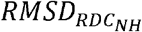 for benchmarking current conformation generation methods, aiming for values close to zero for all. *ε*_*Rg*_ reflects the reasonableness of the model’s generated conformations in terms of their compactness, while *RMSD*s on other physical quantities indicate how well the generated conformations conform to experimental observations on local structures.

For evaluating the conformational ensembles generated by the models, all methods except AF-cluster generated 300 conformations for subsequent evaluation^53,54^. This number of conformations has been previously shown to adequately reflect the structural diversity for intrinsically disordered proteins of similar size to the largest protein in our test set. Additionally, we conducted convergence test on IDPFold-generated ensembles as depicted in Figure S16. The number of conformations generated by AF-cluster is influenced by the number of clusters in the MSA, and we used its default MSA and clustering settings. All conformation generation was performed on NVIDIA A100 GPUs.

## Acknowledgements

This work was supported by the Center for HPC at Shanghai Jiao Tong University, and the National Key Research and Development Program of China (2020YFA0907700 and 2023YFF1205102), the Fundamental Research Funds for the Central Universities (YG2023LC03), and the National Natural Science Foundation of China (21977068, 32171242 and T2222012).

## Code Availability

The code of IDPFold is available at https://github.com/Junjie-Zhu/IDPFold.

