## Supplemental data for "Precise Generation of Conformational Ensembles for Intrinsically Disordered Proteins *via* Fine-tuned Diffusion Models"

**Notes**

The authors declare that there is no conflict of interest.

**Author Contributions**

^#^These authors contributed equally to this work.

**Table S1.** IDP systems in test set, $R_{g}^{exp}$, experimental chemical shifts, J-couplings and RDCs. Abbreviated forms of long system names are marked in bold.

| **System** | $\boldsymbol{R}_{\boldsymbol{g}}^{\boldsymbol{exp}}$ | **Chemical Shifts** | **J-couplings and RDCs** |
| --- | --- | --- | --- |
| Sic1 | 34.7^1^ | $C_{\alpha}$, $C_{\beta}$ chemical shifts^1^ | N/A |
| Aβ 42 | 12.4^2^ | $C_{\alpha}$, $C_{\beta}$ chemical shifts^3^ | ${}^{3}{J(H_{N},H_{\alpha})}$ scalar coupling^4^, backbone N-HN RDCs^5^ |
| Aβ 40 | 12^6^ | $C_{\alpha}$, $C_{\beta}$ chemical shifts^3^ | ${}^{3}{J(H_{N},H_{\alpha})}$ scalar coupling^4^, backbone N-HN RDCs^5^ |
| RS1 | 12.62^7^ | $C_{\alpha}$ chemical shifts^8^ | ${}^{3}{J(H_{N},H_{\alpha})}$ scalar coupling^8^, backbone N-HN RDCs^8^ |
| Histatin5 | 13.8^9^ | N/A | ${}^{3}{J(H_{N},H_{\alpha})}$ scalar coupling^10^ |
| p15^PAF^ | 28.1^11^ | $C_{\alpha}$, $C_{\beta}$ chemical shifts^11^ | backbone N-HN RDCs^11^ |
| FEZ1 monomer | 36^12^ | N/A | N/A |
| HIV-1 Tat_133_ | 33^13^ | N/A | N/A |
| R17 | 22.9^14^ | N/A | N/A |
| PIR domain | 26.5^15^ | N/A | N/A |
| IB5 | 27.9^16^ | N/A | N/A |
| PaaA2 | 22.4^17^ | $C_{\alpha}$, $C_{\beta}$ chemical shifts^17^ | backbone N-HN RDCs^17^ |
| ACTR | 25^18^ | $C_{\alpha}$, $C_{\beta}$ chemical shifts^19^ | backbone N-HN RDCs^20^ |
| drkN SH3 | 16.7^21^ | $C_{\alpha}$, $C_{\beta}$ chemical shifts^22^ | ${}^{3}{J(H_{N},H_{\alpha})}$ scalar coupling^7^ |
| Juxtanodin | 55.9^23^ | N/A | N/A |
| α Synuclein | 33^24^ | $C_{\alpha}$, $C_{\beta}$ chemical shifts^19^ | ${}^{3}{J(H_{N},H_{\alpha})}$ scalar coupling^25^ |
| β Synuclein | 49^26^ | N/A | N/A |
| γ Synuclein | 61^26^ | N/A | N/A |
| Tau K18 | 38^27^ | N/A | N/A |
| Tau K19 | 35^27^ | N/A | N/A |
| Prothymosin α | 37.8^28^ | N/A | N/A |
| p53 (1-93) **[p53]** | 28.7^29^ | $C_{\alpha}$, $C_{\beta}$ chemical shifts^29^ | N/A |
| Human NCBD domain **[NCBD]** | 33^30^ | N/A | N/A |
| Human Calpastatin (137-237) **[Calpastatin]** | 39^30^ | N/A | N/A |
| N-term NRG1 type III **[NtermNRG1]** | 26.8^31^ | N/A | N/A |
| N-term VS Virus phosphoprotein  **[NtermVS]** | 26^32^ | N/A | N/A |
| E3 ubiquitin ligase RNF4 (32-82)  **[RNF4]** | 25.8^33^ | N/A | N/A |

**Table S2.** Sequences of IDPs in test set.

| **System** | **Sequence** |
| --- | --- |
| Sic1 | MTPSTPPRSRGTRYLAQPSGNTSSSALMQGQKTPQKPSQNLVPVTPSTTKSFKNAPLLAPPNSNMGMTSPFNGLTSPQRSPFPKSSVKRTLF |
| Aβ 42 | DAEFRHDSGYEVHHQKLVFFAEDVGSNKGAIIGLMVGGVVIA |
| Aβ 40 | DAEFRHDSGYEVHHQKLVFFAEDVGSNKGAIIGLMVGGVV |
| RS1 | GAMGPSYGRSRSRSRSRSRSRSRS |
| Histatin5 | DSHAKRHHGYKRKFHEKHHSHRGY |
| p15^PAF^ | VRTKADSVPGTYRKVVAARAPRKVLGSSTSATNSTSVSSRKAENKYAGGNPVCVRPTPKWQKGIGEFFRLSPKDSEKENQIPEEAGSSGLGKAKRKACPLQPDHTNDEKE |
| FEZ1 monomer | QIQEEEETLQDEEVWDALTDNYIPSLSEDWRDPNIEALNGNCSDTEIHEKEEEEFNEKSENDSGINEEPLLTADQVIEEIEEMMQNSPDPEEEEEVLEEEDGG |
| HIV-1 Tat_133_ | MEPVDPRLEPWKHPGSQPRTACTNCYCKKCCFHCQVCFIRKALGISYGRKKRRQRRRAPPDSETHQVSPPKQPASQPRGDPTGPKESKKKVERETETHPVN |
| R17 | RLEESLEYQQFVANVEEEEAWINEKMTLVASEDYGDTLAAIQGLLKKHEAFETDFTVHKDRVNDVAANGEDLIKKNNHHVENITAKMKGLKGKVSDLEKA |
| PIR domain | SVSPMRSVSENSLVAMDFSGQKTRVIDNPTEALSVAVEEGLAWRKKGCLRLGNHGSPTAPSQSSAVNMALHRSQP |
| IB5 | SARSPPGKPQGPPQQEGNKPQGPPPPGKPQGPPPAGGNPQQPQAPPAGKPQGPPPPPQGGRPPRPAQGQQPPQ |
| PaaA2 | MDYKDDDDKNRALSPMVSEFETIEQENSYNEWLRAKVATSLADPRPAIPHDEVERRMAERFAKMRKERSKQ |
| ACTR | GTQNRPLLRNSLDDLVGPPSNLEGQSDERALLDQLHTLLSNTDATGLEEIDRALGIPELVNQGQALEPKQD |
| drkN SH3 | MEAIAKHDFSATADDELSFRKTQILKILNMEDDSNWYRAELDGKEGLIPSNYIEMKNHD |
| Juxtanodin | MTDTPVTLSGSECNGDRPPENGQQPSSQTRKTTDADETQTYYGVEPSLQHLPAKENQEESGNSKGNVLPRGSEDEKILNENTEENLFVVHQAIQDLSLQETSAEDTVFQEGHPWKKIPLNSHNLDMSRQKERIVHQHLEQREDESAAHQATEIEWLGFQKSSQVDILHSKCDEEEEVWNEEINEEDVDECAEDEGEDEVRVIEFKRKYREGSPLKEESLAREDSPLSSPSSQPGTPDEQLVLGKKGDIARNSYSRYNTISYRKIRKGNTKQRIDEFESMMHL |
| α Synuclein | MDVFMKGLSKAKEGVVAAAEKTKQGVAEAAGKTKEGVLYVGSKTKEGVVHGVATVAEKTKEQVTNVGGAVVTGVTAVAQKTVEGAGSIAAATGFVKKDQLGKNEEGAPQEGILEDMPVDPDNEAYEMPSEEGYQDYEPEA |
| β Synuclein | MDVFMKGLSMAKEGVVAAAEKTKQGVTEAAEKTKEGVLYVGSKTREGVVQGVASVAEKTKEQASHLGGAVFSGAGNIAAATGLVKREEFPTDLKPEEVAQEAAEEPLIEPLMEPEGESYEDPPQEEYQEYEPEA |
| γ Synuclein | MDVFKKGFSIAKEGVVGAVEKTKQGVTEAAEKTKEGVMYVGAKTKENVVQSVTSVAEKTKEQANAVSEAVVSSVNTVATKTVEEAENIAVTSGVVRKEDLRPSAPQQEGEASKEKEEVAEEAQSGGD |
| Tau K18 | QTAPVPMPDLKNVKSKIGSTENLKHQPGGGKVQIINKKLDLSNVQSKCGSKDNIKHVPGGGSVQIVYKPVDLSKVTSKCGSLGNIHHKPGGGQVEVKSEKLDFKDRVQSKIGSLDNITHVPGGGNKKIE |
| Tau K19 | QTAPVPMPDLKNVKSKIGSTENLKHQPGGGKVQIVYKPVDLSKVTSKCGSLGNIHHKPGGGQVEVKSEKLDFKDRVQSKIGSLDNITHVPGGGNKKIE |
| Prothymosin α | MSDAAVDTSSEITTKDLKEKKEVVEEAENGRDAPANGNAENEENGEQEADNEVDEEEEEGGEEEEEEEEGDGEEEDGDEDEEAESATGKRAAEDDEDDDVDTKKQKTDEDD |
| p53 (1-93) | MEEPQSDPSVEPPLSQETFSDLWKLLPENNVLSPLPSQAMDDLMLSPDDIEQWFTEDPGPDEAPRMPEAAPPVAPAPAAPTPAAPAPAPSWPL |
| Human NCBD domain | ISPLKPGTVSQQALQNLLRTLRSPSSPLQQQQVLSILHANPQLLAAFIKQRAAKYANSNPQPIPGQPGMPQGQPGLQPPTMPGQQGVHSNPAMQNMNPMQAGVQR |
| Human Calpastatin (137-237) | AVPVESKPDKPSGKSGMDAALDDLIDTLGGPEETEEENTTYTGPEVSDPMSSTYIEELGKREVTIPPKYRELLAKKEGITGPPADSSKPIGPDDAIDALSSDFTCGSPTAAGKKTEKEESTEVLKAQSAGTVRSAAPPQEK |
| N-term NRG1 type III | MEIYSPDMSEVAAERSSSPSTQLSADPSLDGLPAAEDMPEPQTEDGRTPGLVGLAV |
| N-term VS Virus phosphoprotein | MDNLTKVREYLKSYSRLDQAVGEIDEIEAQRAEKSNYELFQEDGVEEHTKPSYFQAADDSHHHHHHHH |
| E3 ubiquitin ligase RNF4 (32-82) | EAEPIELVETAGDEIVDLTCESLEPVVVDLTHNDSVVIVDERRRPRRNARR |

**Table S3.** Performance of all 8 versions of AlphaFlow.

| **Methods** | **Validity (↑)** | **Fidelity** | |
| --- | --- | --- | --- |
|  |  | $\varepsilon_{Rg}$ | ${MAE}_{\delta C_{\alpha}}$ (↓) |
| AF-PDB-base | **0.97** | $\boldsymbol{-0.24}$ | **0.54** |
| AF-PDB-distilled | 0.91 | $-0.33$ | 0.78 |
| AF-MD-base | 0.94 | $-0.32$ | **0.54** |
| AF-MD-distilled | 0.91 | $-0.44$ | 0.66 |
| ESM-PDB-base | 0.94 | $\boldsymbol{-0.24}$ | 0.64 |
| ESM-PDB-distilled | 0.89 | $-0.33$ | 0.77 |
| ESM-MD-base | 0.96 | $-0.35$ | 0.62 |
| ESM-MD-distilled | 0.90 | $-0.47$ | 0.68 |

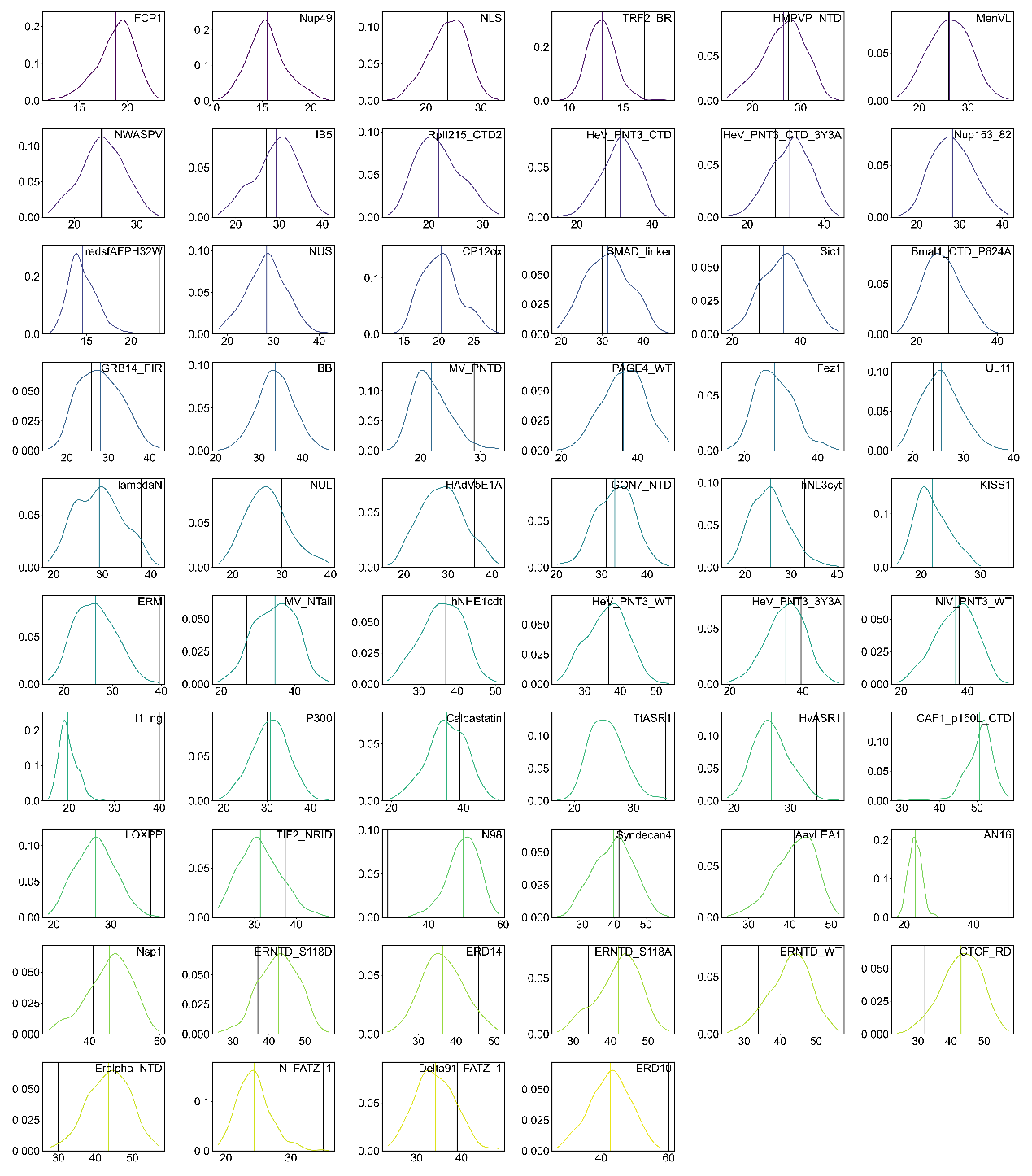

**Figure S1.** Rg distribution of IDPFold generated conformation ensembles (colored) on 58 IDP systems from IDRome. Experimental Rg values are plotted as black solid lines.

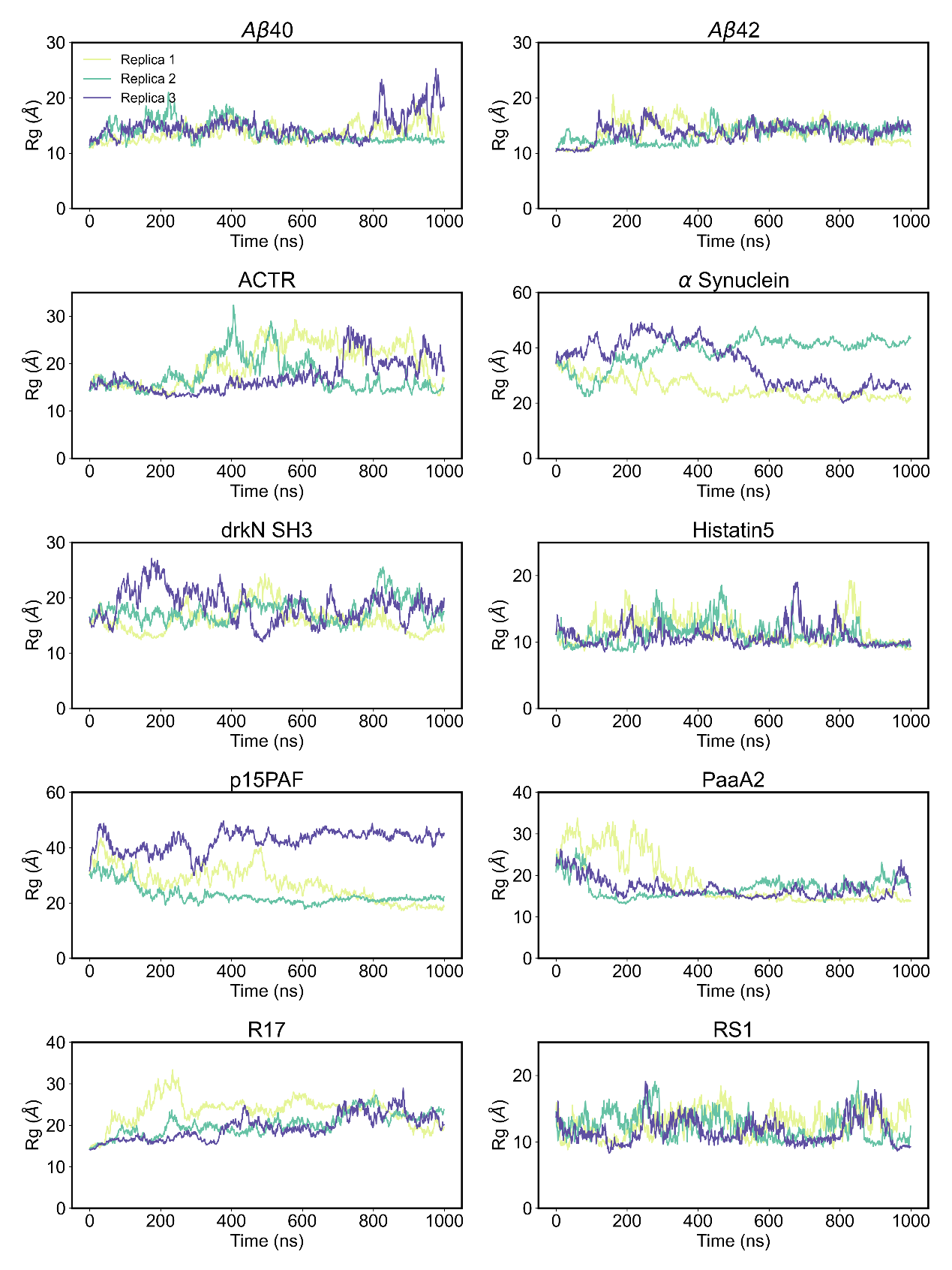

**Figure S2.** Convergence of all-atom MD simulation.

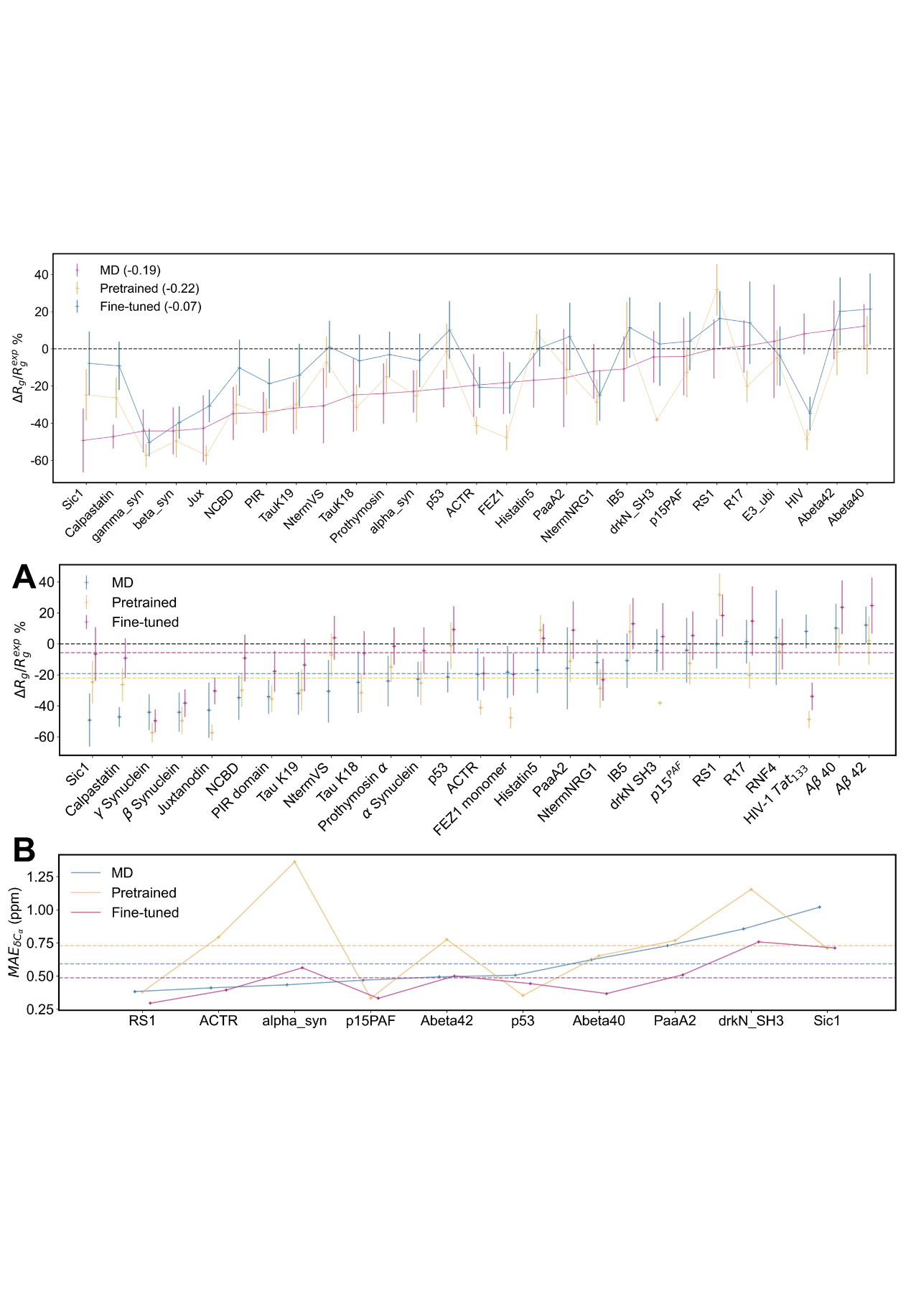

**Figure S3.** Differences between generated conformational ensembles and experimental observations. (A) Average Rg errors and their standard deviation on all 27 IDPs. (B) Mean average error on ensemble average $C_{\alpha}$ chemical shift on all 10 IDPs.

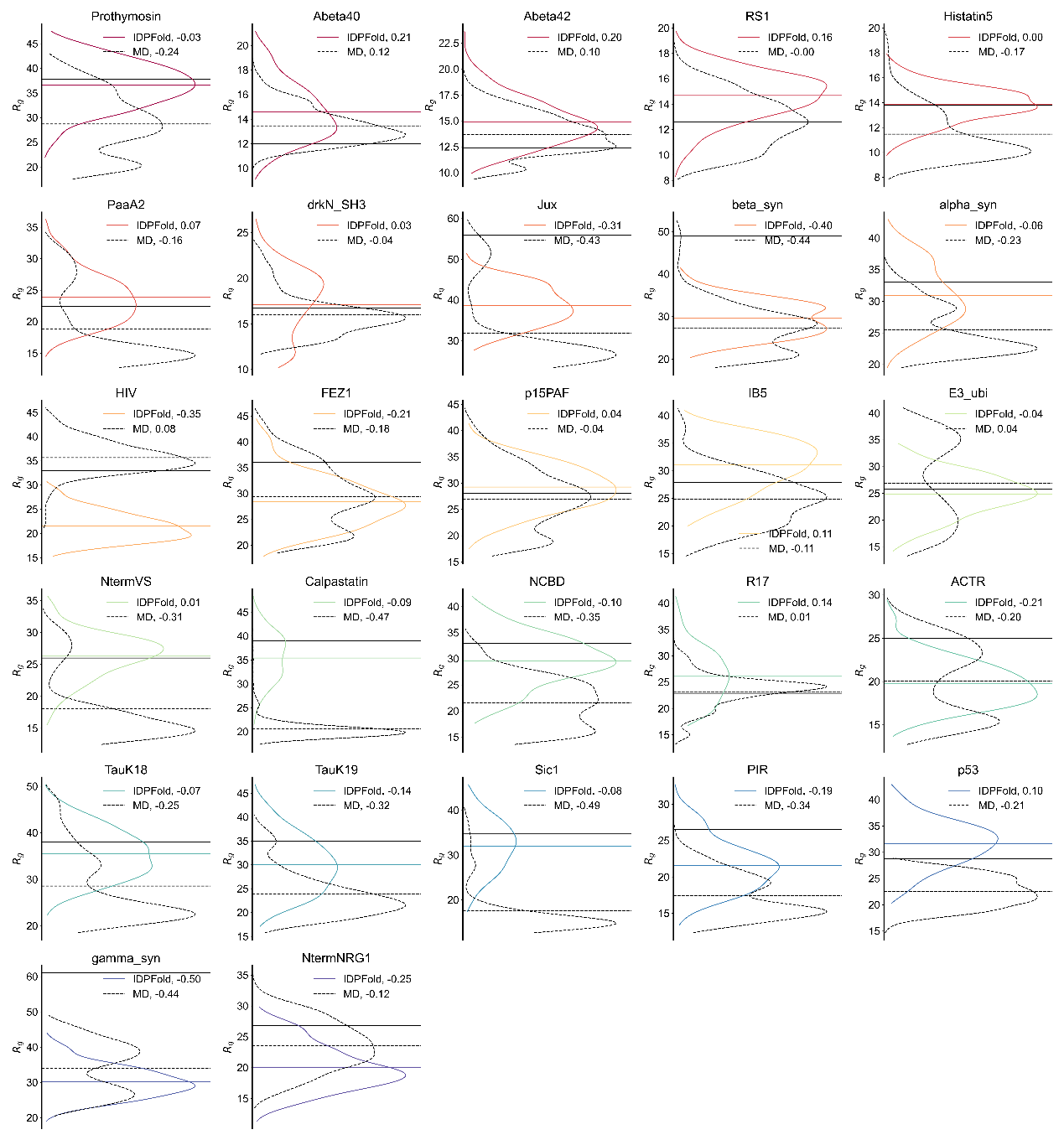

**Figure S4.** Rg distribution of **IDPFold** generated conformation ensembles (colored) and MD trajectories (black dashed) on 27 IDP systems. Experimental Rg values are plotted as black solid lines.

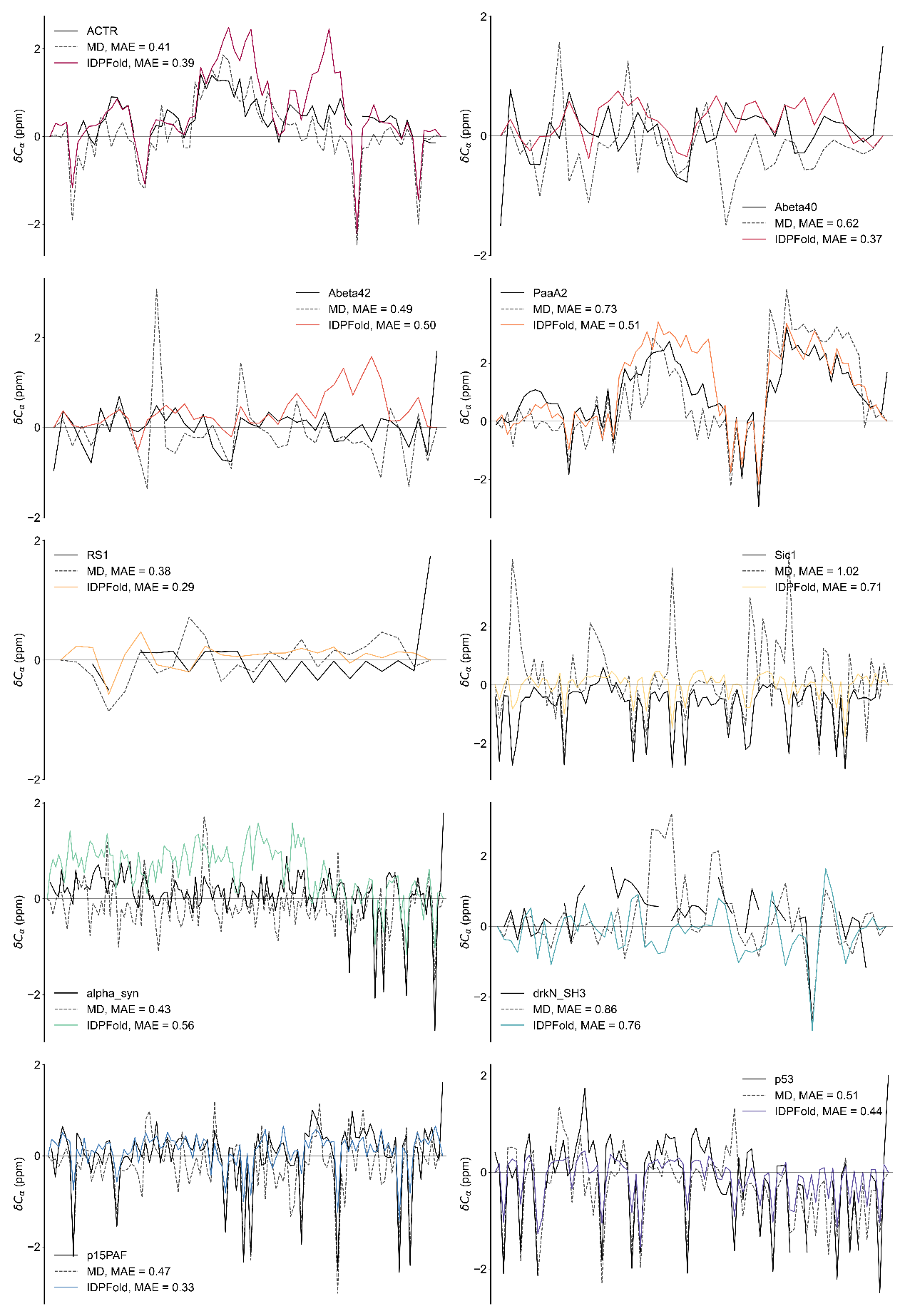

**Figure S5.** Ensemble average $C_{\alpha}$ chemical shifts of **IDPFold** generated conformation ensembles (colored) and MD trajectories (black dashed) on 10 IDP systems. Experimental values are plotted as black solid lines.

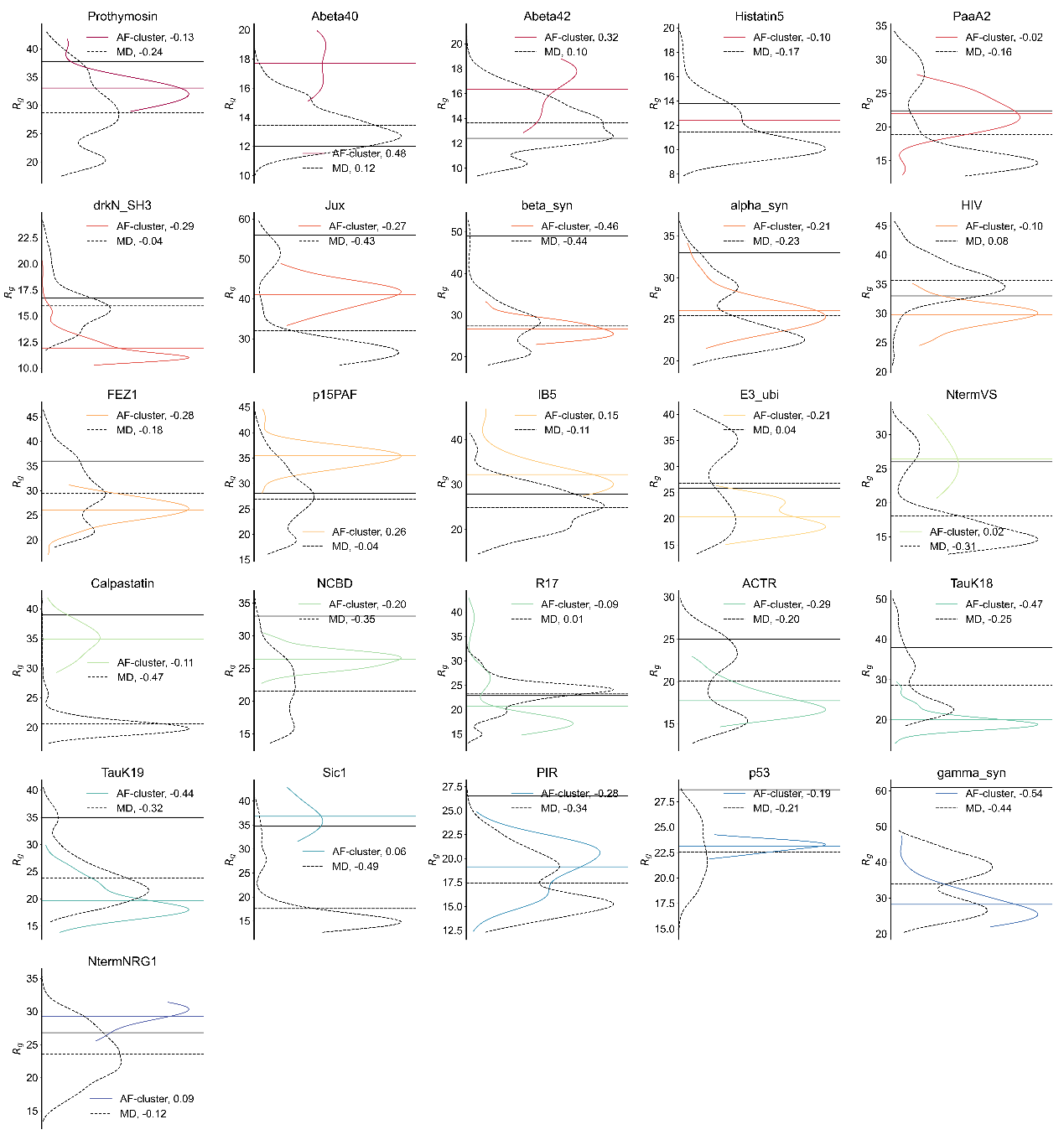

**Figure S6.** Rg distribution of **AF-cluster** generated conformation ensembles (colored) on 26 IDP systems. Experimental Rg values are plotted as black solid lines.

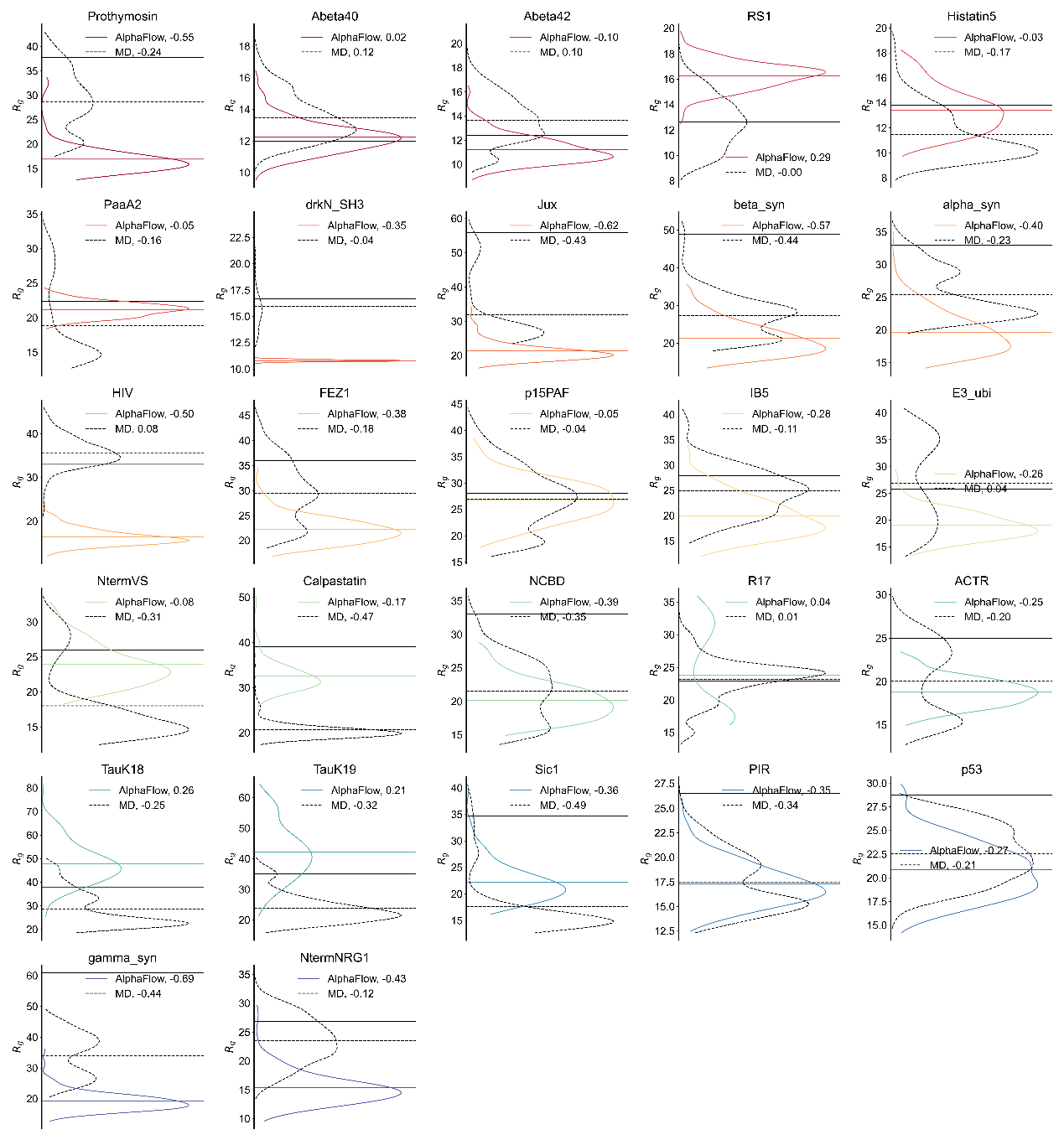

**Figure S7.** Rg distribution of **AlphaFlow** (PDB-base) generated conformation ensembles (colored) on 27 IDP systems. Experimental Rg values are plotted as black solid lines.

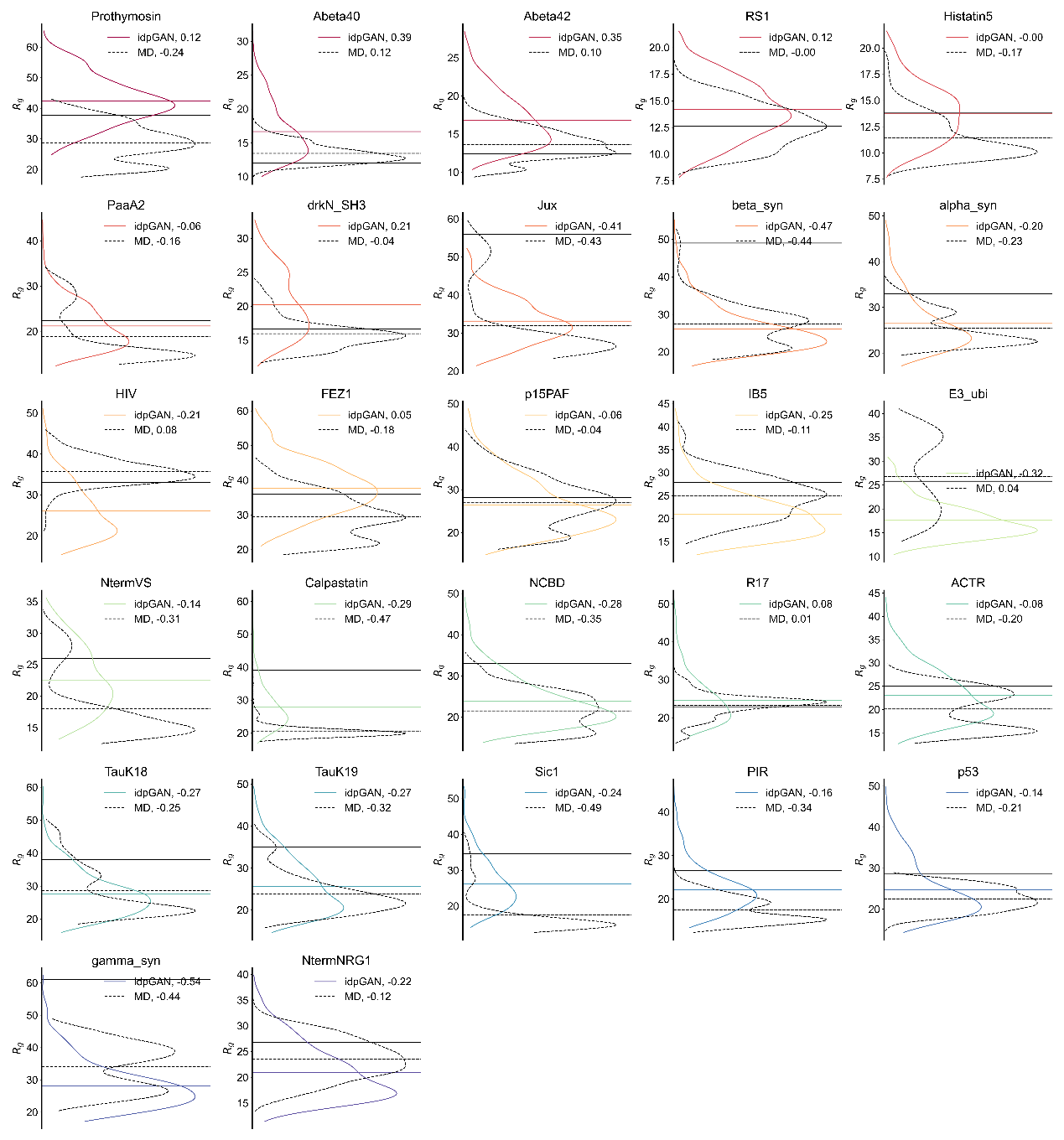

**Figure S8.** Rg distribution of **idpGAN** generated conformation ensembles (colored) on 27 IDP systems. Experimental Rg values are plotted as black solid lines.

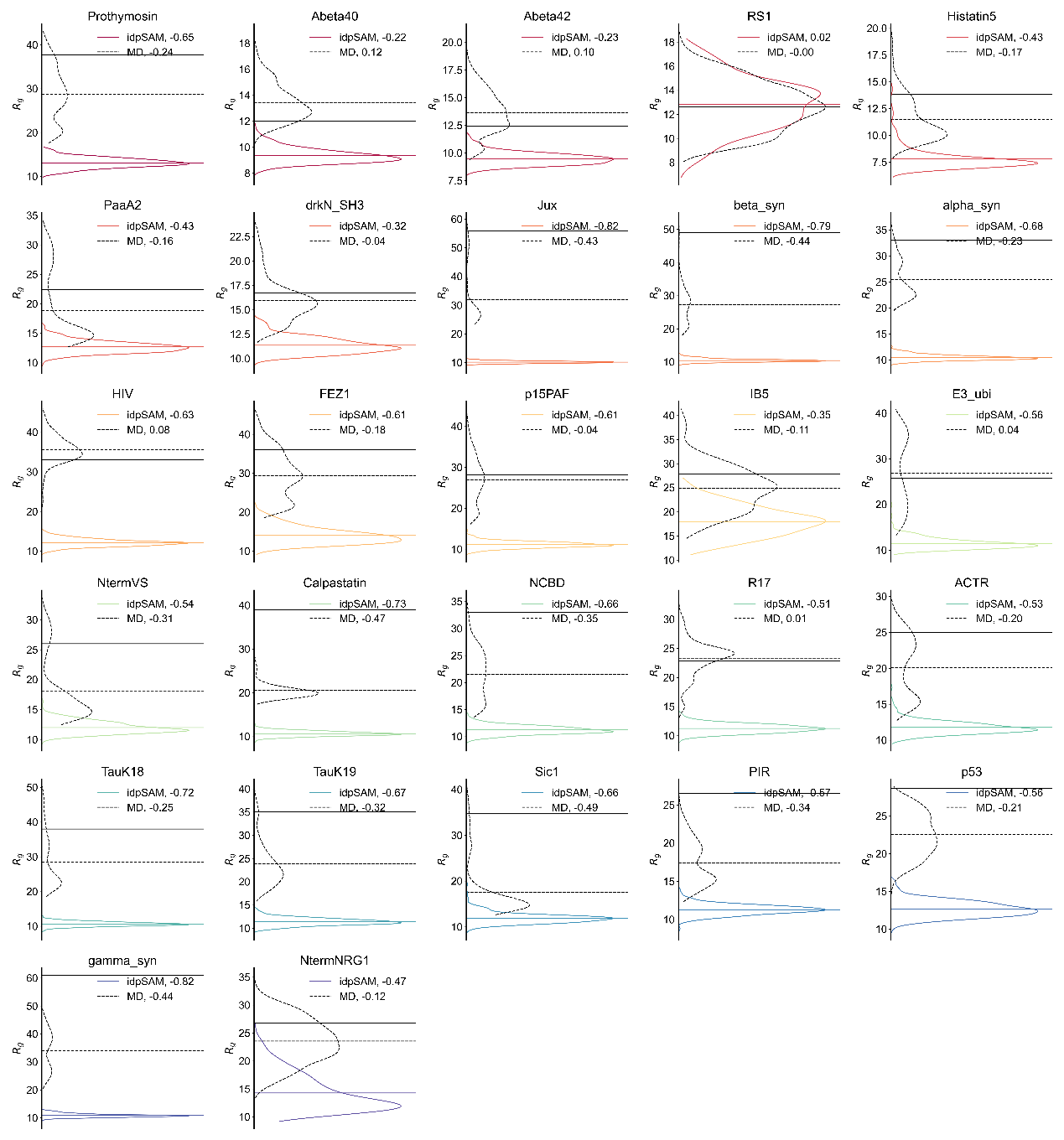

**Figure S9.** Rg distribution of **idpSAM** generated conformation ensembles (colored) on 27 IDP systems. Experimental Rg values are plotted as black solid lines.

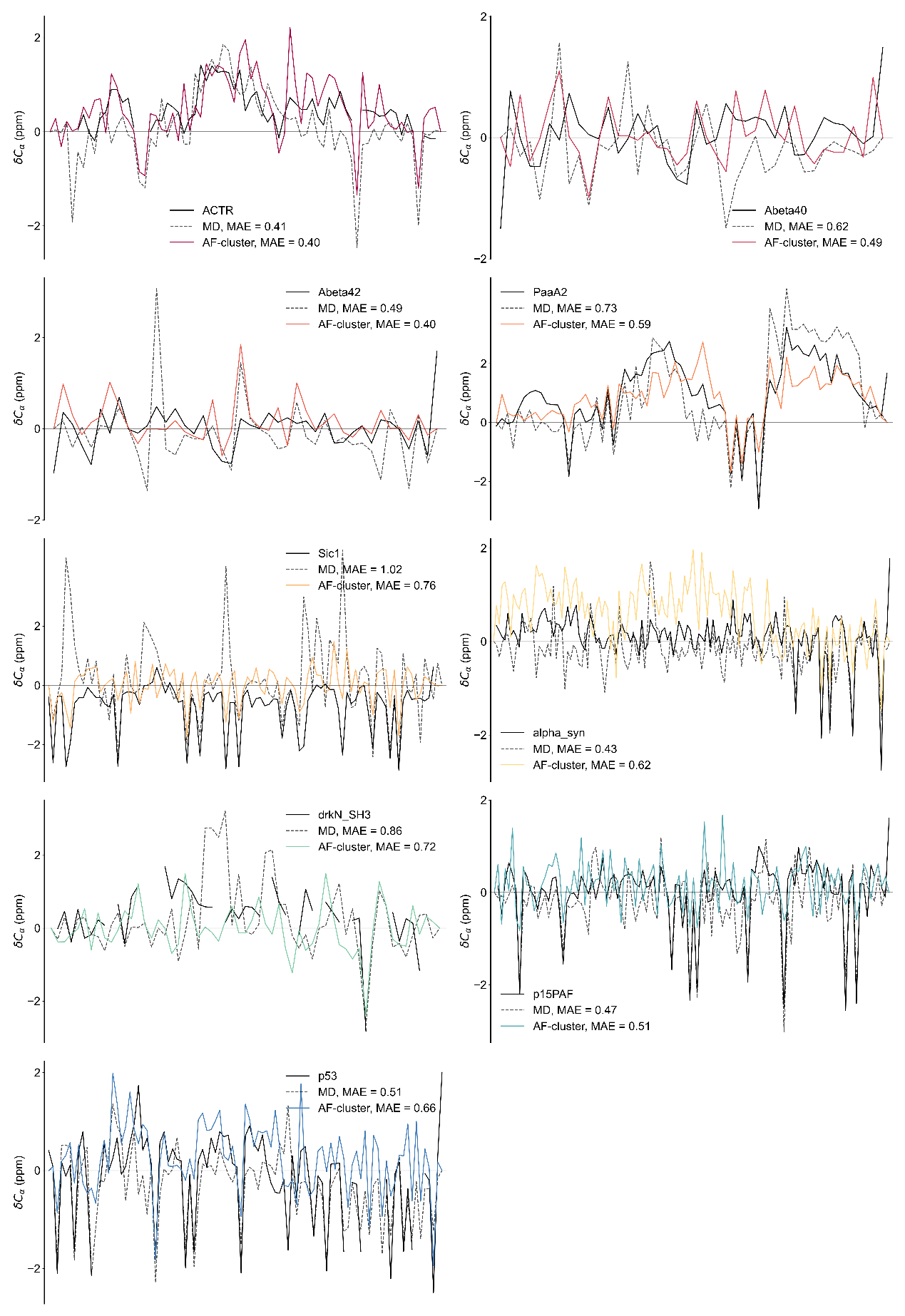

**Figure S10.** Ensemble average $C_{\alpha}$ chemical shifts of **AF-cluster** generated conformation ensembles (colored) on 10 IDP systems. Experimental values are plotted as black solid lines.

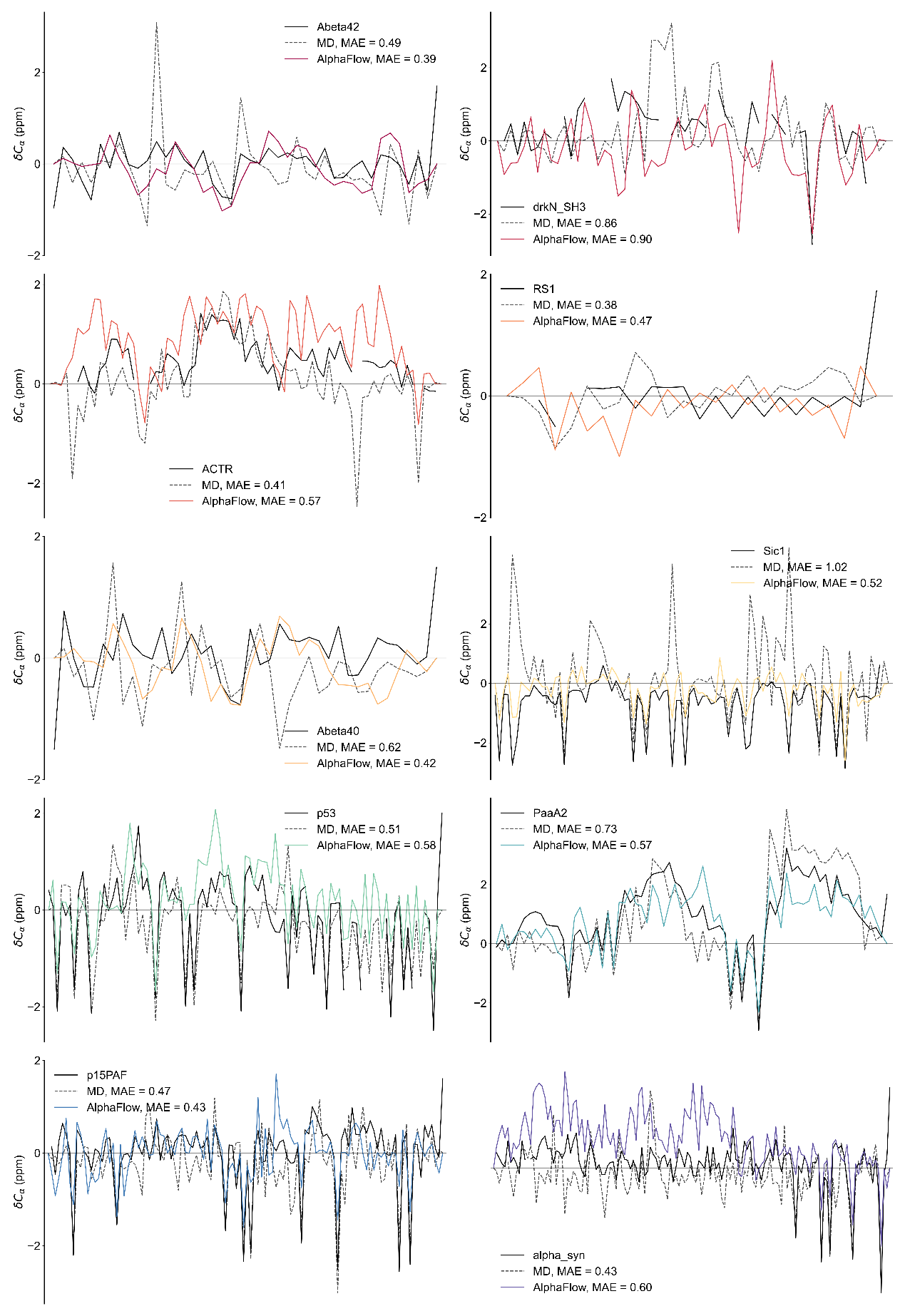

**Figure S11.** Ensemble average $C_{\alpha}$ chemical shifts of **AlphaFlow** (PDB-base) generated conformation ensembles (colored) on 10 IDP systems. Experimental values are plotted as black solid lines.

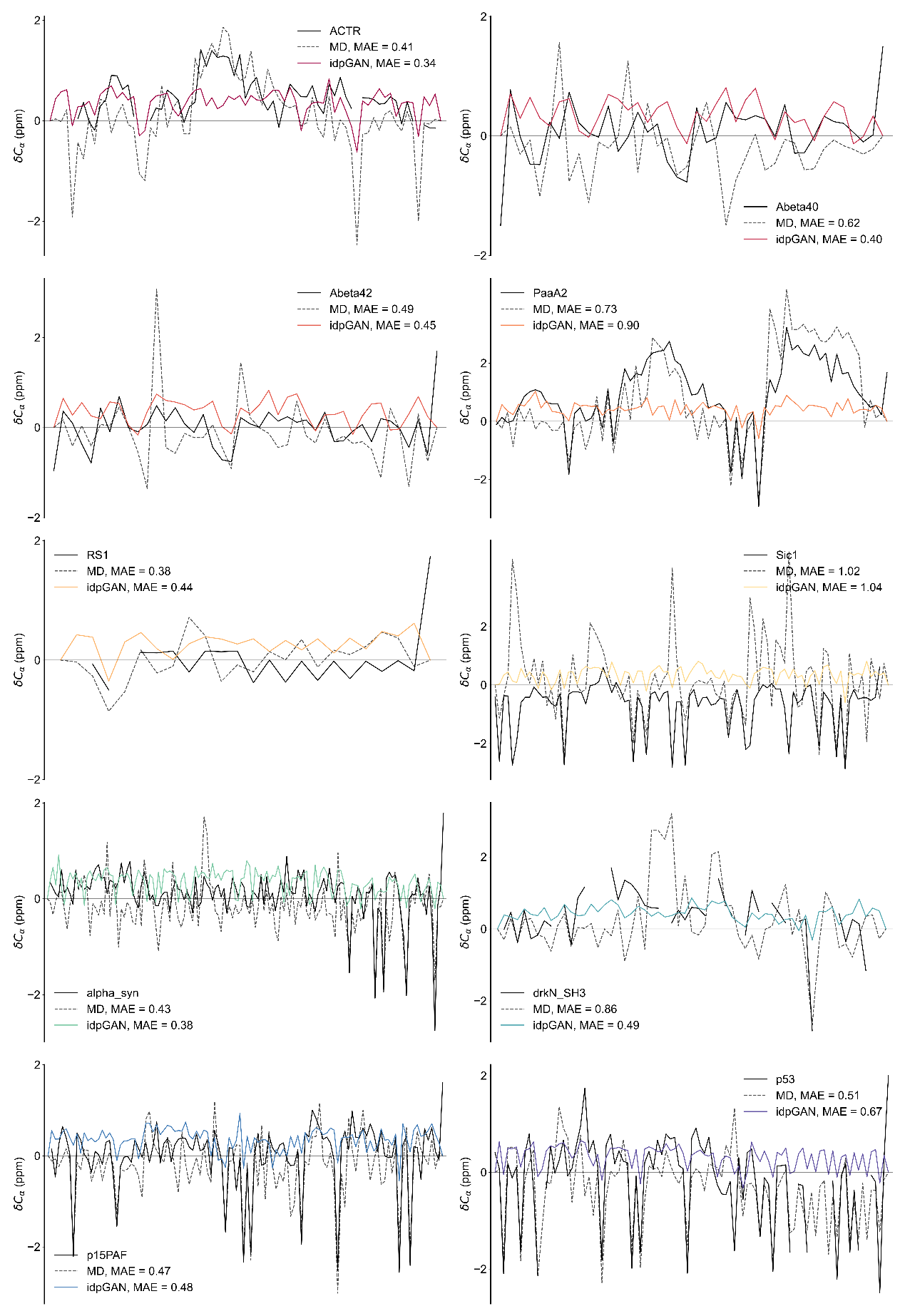

**Figure S12.** Ensemble average $C_{\alpha}$ chemical shifts of force-field-refined **idpGAN** generated conformation ensembles (colored) on 10 IDP systems. Experimental values are plotted as black solid lines.

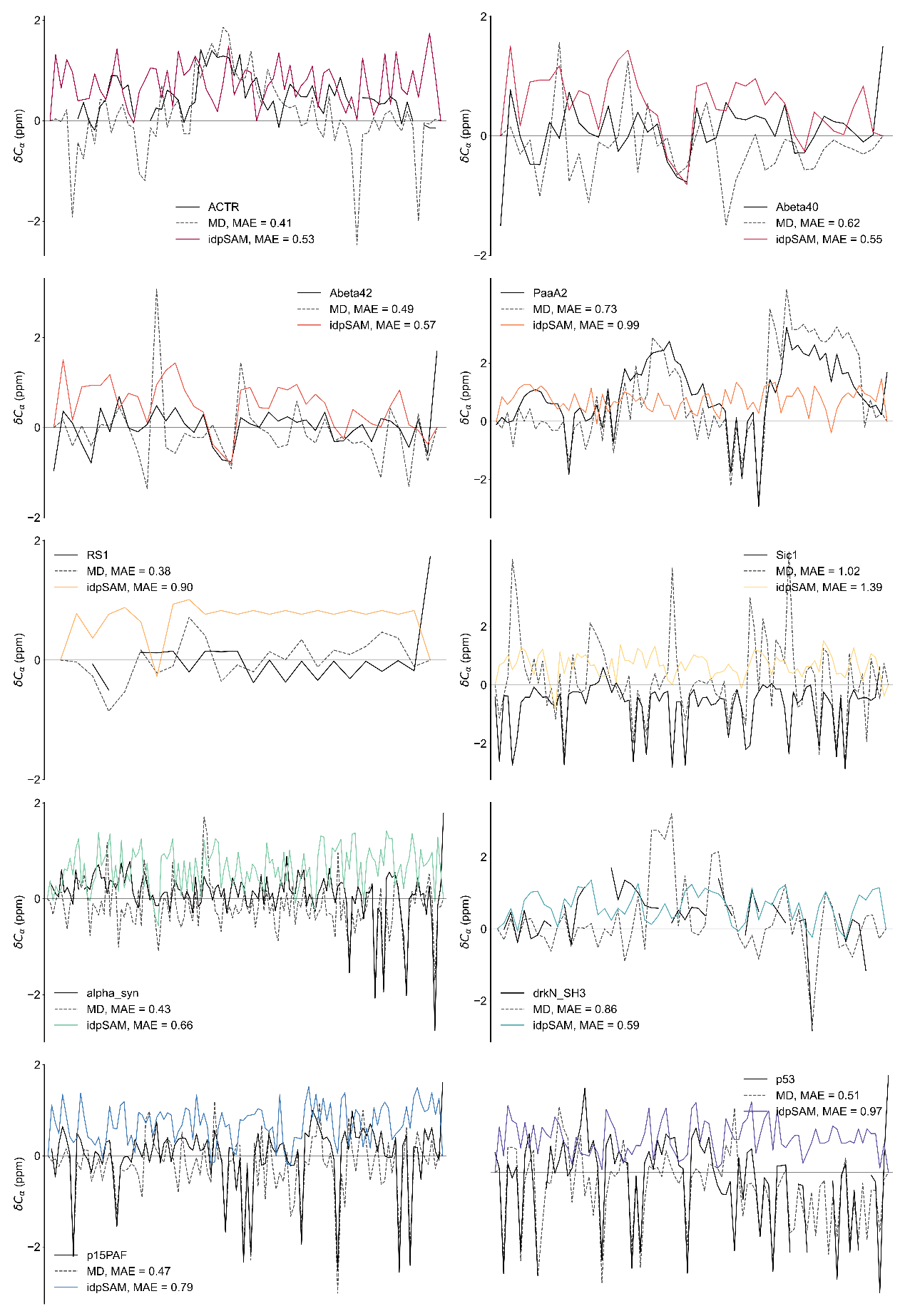

**Figure S13.** Ensemble average $C_{\alpha}$ chemical shifts of force-field-refined **idpSAM** generated conformation ensembles (colored) on 10 IDP systems. Experimental values are plotted as black solid lines.

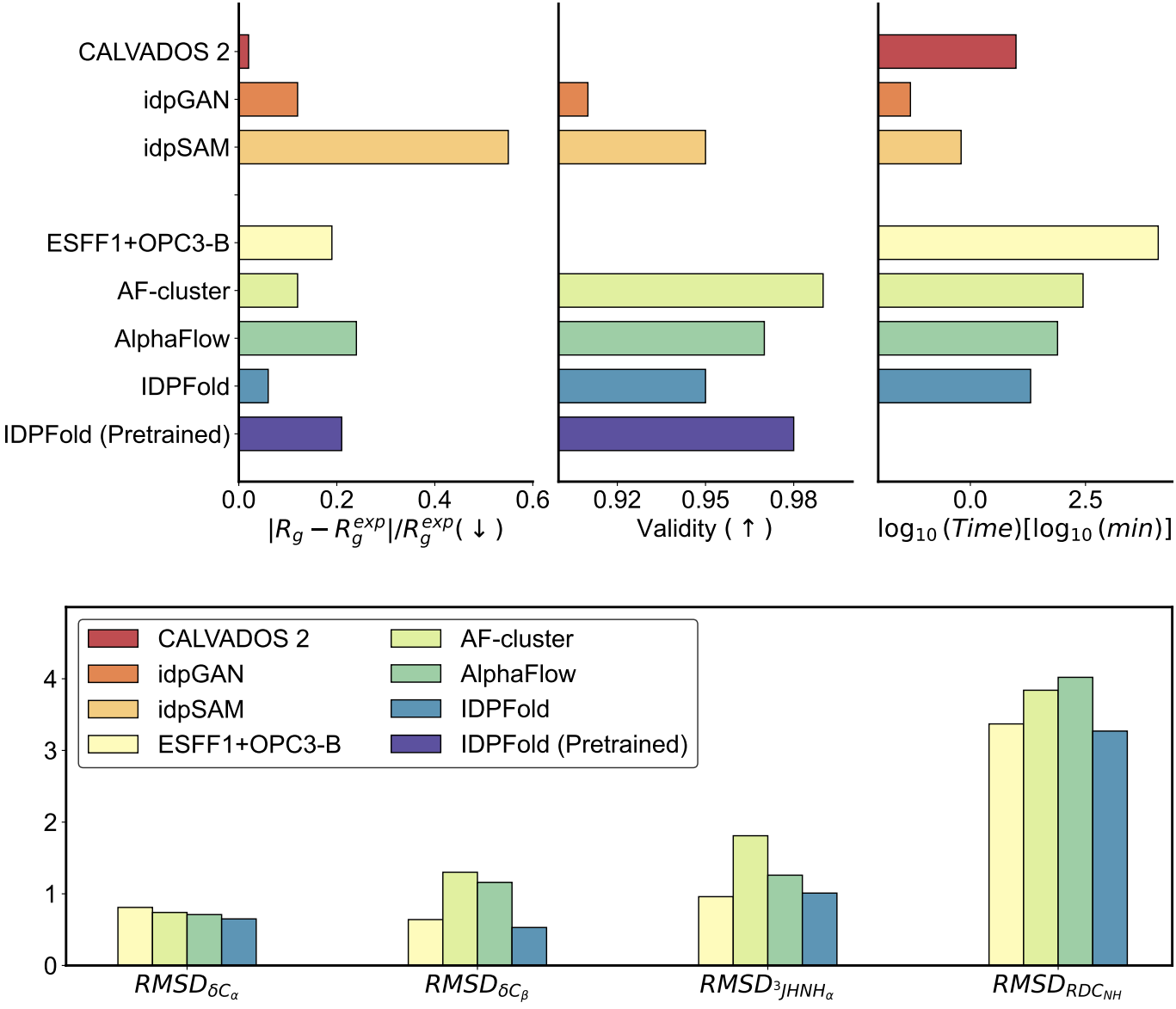

**Figure S14.** Benchmark of generative deep learning methods on IDP ensemble prediction.

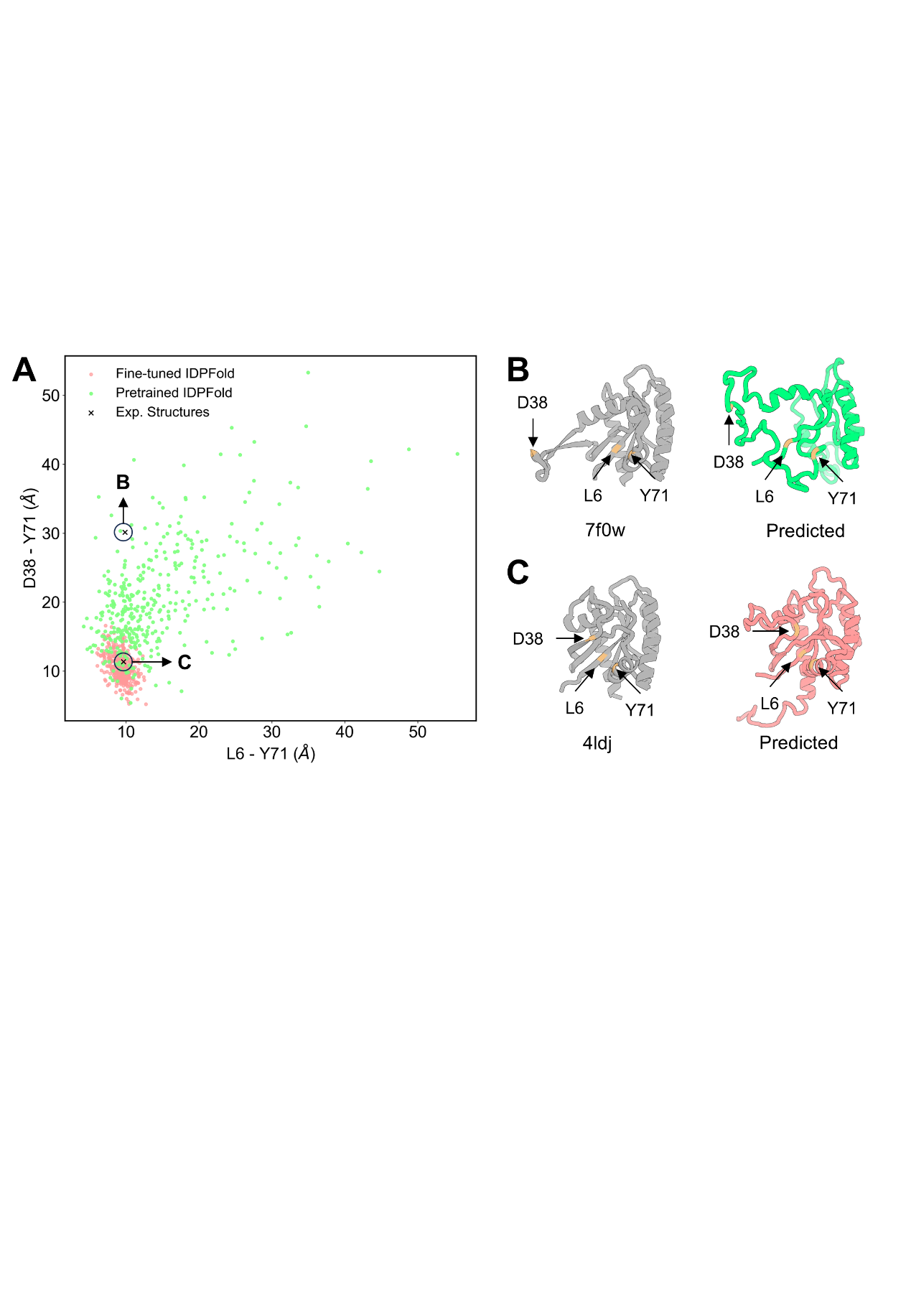

**Figure S15.** IDPFold generates a wide range of conformations for allosteric proteins. (A) IDPFold generated conformations mapped on L6-Y71-D38 distance plane. (B) Fine-tuned IDPFold generated conformation compared to X-ray structure (PDB ID: 7f0w). (C) Pretrained IDPFold generated conformation compared to X-ray structure (PDB ID: 4ldj)^34^.

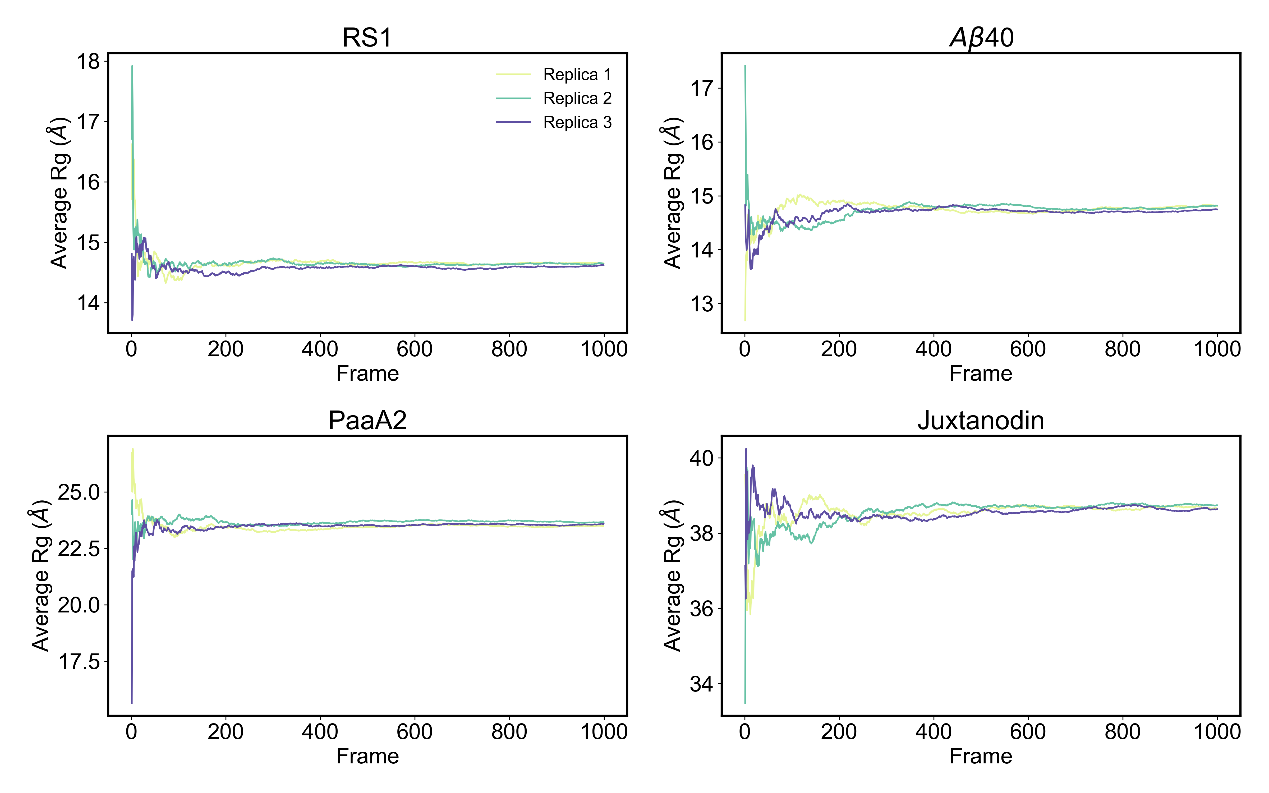

**Figure S16.** Convergence of IDPFold-generated ensembles.
